# Calcium directs actin assembly via allosteric activation of formin INF2

**DOI:** 10.64898/2026.01.27.701936

**Authors:** Bohan Zhang, Meng Zhang, Ke Liu, Chenhui Zhao, Jianwen Zhang, Zhijun Liu, Ying Fan, Ruo-xu Gu, Lin Lin, Chuanhai Fu, Jinwei Zhu

## Abstract

Calcium signals spatiotemporally orchestrate cytoskeletal dynamics, typically through Rho GTPase-mediated signaling cascades, yet the direct molecular transducers that convert local calcium transients into spatially controlled actin assembly have remained elusive. Here, we uncover a structure-based mechanism by which calcium-bound calmodulin (Ca^2+^-CaM) directly activates formin INF2 to drive actin assembly. We demonstrate that Ca^2+^-CaM binds to the diaphanous inhibitory domain (DID) of INF2 with nanomolar affinity, inducing allosteric conformational changes that sterically disrupt INF2 autoinhibition. The unexpected bipartite Ca^2+^-CaM binding interface on INF2 enables ultrasensitive decoding of local calcium microdomains, such as ER-mitochondrial contacts, where activated INF2 promotes actin polymerization to facilitate mitochondrial fission. We further show that the Charcot-Marie-Tooth disease–associated INF2 R91G mutation enhances Ca^2+^-CaM binding via optimized interfacial dynamics, suggesting a gain-of-function disease mechanism. Our work establishes the Ca^2+^-CaM–INF2 axis as a direct molecular transducer linking spatial calcium signals to actin-dependent organelle dynamics, defining a Rho-GTPase-independent paradigm for calcium–cytoskeleton communication, with broad implications for INF2-linked pathologies.

## Introduction

Actin filament assembly dynamically shapes cellular architecture and drives cell morphogenesis in eukaryotes ^1^. Formins, a family of actin-binding proteins, regulate actin assembly in fundamental cellular processes such as cell division, cell motility, organelle dynamics and tissue morphogenesis ^2,3^. Formins possess conserved formin homology 1 (FH1) and FH2 domains, which synergistically mediate nucleation and processive elongation of actin filaments ^4^. These catalytic modules are often flanked by divergent regulatory domains, including the N-terminal Rho-binding domain (RBD) and diaphanous inhibitory domain (DID), and the C-terminal diaphanous autoregulatory domain (DAD) ^2^. The activity of these diaphanous-related formins (DRFs) must be precisely controlled to orchestrate spatiotemporally actin cytoskeleton dynamics. Most DRFs, such as mammalian diaphanous 1 (mDia1), are autoinhibited in their basal state. An intramolecular DID–DAD interaction sterically occludes the FH1-FH2 catalytic core, thereby preventing actin polymerization ^5,6^. Binding of an active Rho GTPase to the RBD disrupts the DID–DAD interaction, resulting in the relief of autoinhibition and formin activation ^7–9^.

Inverted formin 2 (INF2) is an atypical DRF with unique structural and functional features ^10^. Notably, it has two splice variants with different C-terminal sequences: INF2-CAAX, which is anchored to the endoplasmic reticulum (ER) via its CAAX motif, and the cytosolic INF2-nonCAAX **(Fig. 1a)** ^11–13^. INF2 can modulate both actin- and microtubule-based cytoskeletons ^14^, with essential roles in regulation of organelle dynamics, cell motility, and gene expression ^15–18^. Mutations in *INF2* are associated with two distinct diseases: the renal disease focal segmental glomerulosclerosis (FSGS) and Charcot-Marie-Tooth (CMT) neuropathy ^19,20^. Like other DRFs, INF2 adopts an autoinhibited state in cells, as evidenced by direct intramolecular interaction between DID and DAD ^11,21^. This autoinhibition is further supported by the observation that mutation of a key DID residue (A149D, analogous to a variant that disrupts DID–DAD binding in mDia1), induces constitutive INF2 activation ^15,17,22^. However, the absence of a canonical RBD in INF2 implies a distinct regulatory mechanism governing INF2 activation.

**Fig. 1.**
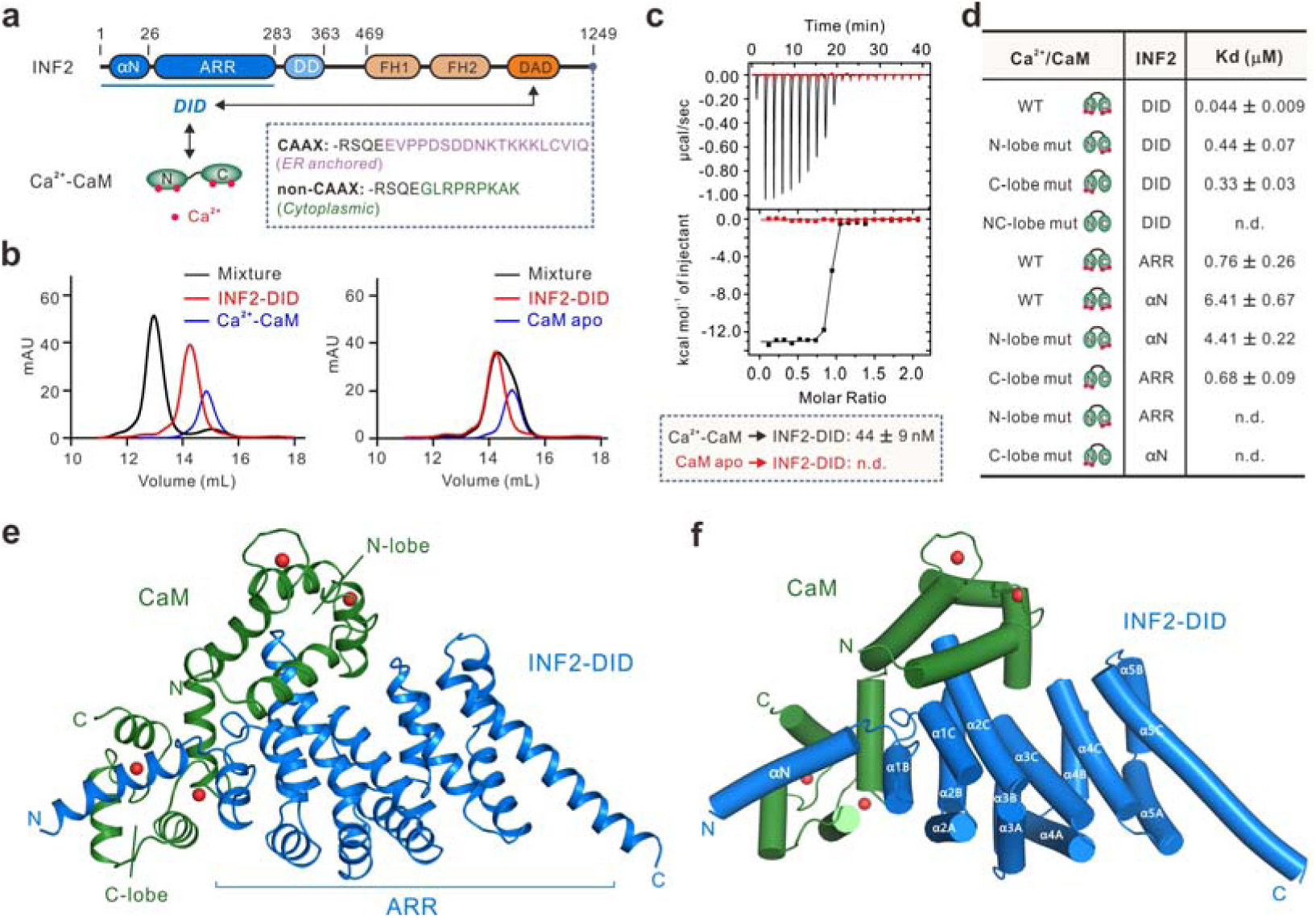
Direct interaction between Ca^2+^-CaM and INF2 DID. **(a)** Schematic diagram of the domain organization of INF2 and Ca^2+^-CaM. The interaction between DID and DAD is indicated by a two-way arrow. Of note, INF2 has two splice variants with different C-terminal sequences: the ER-bound INF2-CAAX and cytosolic INF2-nonCAAX. DID, diaphanous inhibitory domain; ARR, armadillo repeats; DD, dimerization domain; DAD, diaphanous autoregulatory domain; FH, formin homology; αN, N-terminal amphipathic α-helix. CaM consists of two lobes (N- and C-lobe), with each lobe binding two Ca^2+^. **(b)** Analytical gel filtration chromatography assays showed that INF2-DID forms a stable complex with Ca^2+^-CaM, but not with Apo-CaM. The concentration of each protein was 50 µM, and assays were performed at 25 °C. **(c)** Isothermal titration calorimetry (ITC) analysis of bindings of INF2 DID (50 μM) to Ca2+-CaM (black) and Apo-CaM (red) (500 μM). n.d., not detectable. **(d)** Summary of ITC-derived binding affinities between WT and mutants of INF2 DID and Ca^2+^-CaM. n.d., not detectable. **(e)** Ribbon diagram of the overall structure of the INF2-DID (blue) in complex with Ca^2+^-CaM (green). Calcium ions are shown as red spheres. **(f)** Cylinder representation of the INF2 DID–Ca^2+^-CaM complex structure, shown in the same orientation as panel **(e)**.

Emerging evidence implicates calcium signaling as a critical modulator of INF2 activity across distinct cellular contexts ^16,17,23–25^. During mitochondrial fission, ER wraps around mitochondria to induce calcium transients at ER-mitochondria contact sites ^26,27^. The ER-anchored INF2-CAAX is then activated to drive rapid actin polymerization around ER, initiating mitochondrial pre-constriction prior to Drp1-mediated mitochondrial fission ^15,28,29^. In addition, calcium-mediated activation of INF2 at the ER has been proposed to facilitate cellular adaptations during processes like cortex repair and wound healing ^17^.

Furthermore, GPCR signaling elicits acute calcium transients that activate INF2 at the inner nuclear membrane, promoting nuclear actin polymerization for rapid chromatin remodeling ^16^. Despite these context-specific regulatory paradigms, very little is known about the molecular mechanisms underlying calcium-dependent activation of INF2.

Here, we uncover a direct, nanomolar-affinity interaction between Ca^2+^-CaM and INF2 DID. Binding of Ca^2+^-CaM activates INF2, thereby driving actin assembly and promoting mitochondrial fission. The high-resolution crystal structure of the Ca^2+^-CaM–INF2 DID complex unveils a unique allosteric mechanism underlying this calcium-dependent INF2 activation, which is distinct from the canonical Rho-GTPase–mediated formin activation. Furthermore, we demonstrate that the Charcot-Marie-Tooth disease-associated INF2 R91G mutation enhances Ca^2+^-CaM binding via optimized interfacial dynamics, providing a structural and mechanistic framework for this gain-of-function pathology. Collectively, our findings establish the Ca^2+^-CaM–INF2 axis as the long-sought molecular link that transduces spatial calcium signals into localized actin assembly, and position it as a promising therapeutic target for INF2-associated disorders, including CMT and FSGS.

## Results

### Direct interaction between Ca^2+^-CaM and INF2 DID

Considering the lack of intrinsic calcium-binding motifs in INF2 itself, calcium-sensor proteins such as CaM were hypothesized to mediate the calcium-mediated activation of INF2 ^17^. To validate the INF2–CaM interaction, we first confirmed that GST-tagged CaM bound robustly to full-length INF2 in a Ca^2+^-dependent manner **(Fig. S1a)**. Truncation mapping revealed that the DID alone (aa 1-283), rather than the FH1-FH2-DAD tandem, was sufficient to bind to Ca^2+^-CaM **(Fig. S1b-S1d)**. Consistently, analytical gel filtration chromatography assays further confirmed direct complex formation between purified INF2 DID and Ca^2+^-CaM **(Fig. 1b)**. Isothermal titration calorimetry (ITC)-based assay showed that INF2 DID binds to Ca^2+^-CaM with a dissociation constant (*K*_d_) of 44 nM, but not to Ca^2+^-free CaM **(Fig. 1c)**. These results highlighted a direct and high-affinity interaction between INF2 DID and Ca^2+^-CaM.

Sequence analysis of INF2 DID reveals it comprises several armadillo repeats (ARR), preceded by an N-terminal amphipathic α-helix (aa 1-26, referred to as αN) which contains a putative CaM-binding motif **(Fig. 1a)** ^30^. We sought to dissect whether INF2 DID interacts with Ca^2+^-CaM via αN motif. ITC data showed that αN binds to Ca^2+^-CaM with a *K*_d_ value of 6.41 μM **(Figs.1d and S2)**, over 100-fold weaker than that of DID, indicating that ARR may also contribute to the tight association. Indeed, we found that ARR binds to Ca^2+^-CaM, with a *K*_d_ value of 0.76 μM **(Figs.1d and S2)**, highlighting a previously uncharacterized interaction mode for both ARR and CaM. These results suggested that INF2 DID contains two distinct binding sites for Ca^2+^-CaM. Analytical gel filtration chromatography coupled with static light scattering assay revealed a 1:1 stoichiometry of the INF2 DID–Ca^2+^-CaM complex **(Fig. S3a)**, implying that αN and ARR of INF2 DID simultaneously engage one molecule of Ca^2+^-CaM.

CaM has two globular lobes (N- and C-lobe) connected by a flexible central linker. Each lobe binds to two Ca^2+^ via two EF hand motifs ^31^. To test whether both lobes of CaM are involved in the interaction, we generated CaM mutations deficient in Ca^2+^-binding: the N-lobe^mut^ (D21A/D57A) and the C-lobe^mut^ (D94A/D130A) ^32^. We found that the N-lobe^mut^ and C-lobe^mut^ mutation of CaM bound to INF2 DID with *K*_d_ values of 0.44 μM and 0.33 μM, respectively, while the quadruple mutant CaM_NC^mut^ (D21A/D57A/D94A/D130A) totally abolished the binding **(Figs.1d and S2)**. These results demonstrated that both lobes of CaM are required for the high-affinity interaction with INF2 DID. To further assess lobe-specific binding preferences, we performed cross-binding assays between CaM lobe mutations and discrete DID subdomains. The N-lobe^mut^ mutant of CaM lost binding to ARR but retained binding to αN, while the C-lobe^mut^ mutant of CaM lost binding to αN but preserved interaction with ARR **(Figs.1d and S2)**. These results indicated that ARR and αN of DID specifically bind to the N- and C-lobe of Ca^2+^-CaM, respectively.

### Crystal structure of INF2 DID in complex with Ca^2+^-CaM

To uncover the molecular mechanism governing the interaction between INF2 DID and Ca^2+^-CaM, we attempted to determine the complex structure. Despite extensive trials, we could not crystallize the complex. However, the complex prepared from a truncated form of INF2 DID carrying a deletion of a 6-residue flexible loop between αN and ARR (DID^del6^, with aa 20-25 deleted, **Fig. S4**) could be successfully crystallized. It is noteworthy that INF2 DID^del6^ displayed comparable affinity to Ca^2+^-CaM as that of wild-type (WT) INF2 DID **(Fig. S2)**. The complex is referred to as the INF2 DID–Ca^2+^-CaM hereafter for simplicity. The complex structure was solved at 2.11-Å resolution **(Table S1)**. Each asymmetric unit contains two copies of the complex with very similar conformation **(Fig. S3b)**.

In the complex structure, Ca^2+^-CaM adopts an extended conformation, tightly engaging INF2 DID with a buried interface area of 2,188 Å^2^ **(Figs. 1e and S3c)**. Consistent with our biochemical data, the two lobes of Ca^2+^-CaM bind to αN and ARR of INF2 DID in an antiparallel manner, with the C-lobe bound to αN and the N-lobe bound to ARR, respectively **(Fig. 1e and 1f)**. The ARR of INF2 is composed of 14 consecutive α-helices assembled into five armadillo repeats **(Figs. 1f and S3d)**. In classic ARR-mediated interactions, armadillo repeats typically form a superhelical concave groove to recognize extended target peptides ^33^. By contrast, INF2 ARR binds to the well-folded N-lobe of Ca^2+^-CaM through a distinct interface which is remote from the canonical target binding groove **(Fig. S3d)**. This structural feature highlights a previously unanticipated target recognition mode for armadillo repeats.

### Detailed interface of the INF2 DID–Ca^2+^-CaM complex

The INF2 DID–Ca^2+^-CaM interface can be naturally divided into two major sites: the C-lobe–αN site and the N-lobe–ARR site **(Fig. 2a)**. In general, the binding of INF2 DID to Ca^2+^-CaM is mainly mediated by extensive hydrophobic interactions, complemented by several polar contacts. Specifically, at the C-lobe–αN site, W11, L14, and L18 from αN_INF2 make hydrophobic contacts with V92, F93, L106, V109, M110, L113, M125, V137, F142, M145, and M146 from the C-lobe_CaM **(Fig. 2b)**. Several electrostatic interactions further reinforce the interaction, such as K4^INF2^–E128^CaM^, K10^INF2^–E121^CaM^, and K17^INF2^–E115^CaM^ interactions **(Fig. 2b)**. At the N-lobe–ARR site, I92, A95, L96, and L99 from ARR_INF2 form hydrophobic interaction networks with L19, F20, I28, L33, V36, L40, V56, I64, F69, M72, and M73 from the N-lobe_CaM **(Fig. 2c)**. The N-lobe–ARR interface is further stabilized by hydrophilic interactions such as R54/K55^INF2^–E15^CaM^ and N49^INF2^–E12^CaM^ pairs **(Fig. 2c)**. Importantly, the residues that contribute to the binding interfaces are highly conserved in INF2 from different species **(Fig. S4)**, implicating indispensable roles of Ca^2+^-dependent INF2–CaM interaction during evolution.

**Fig. 2.**
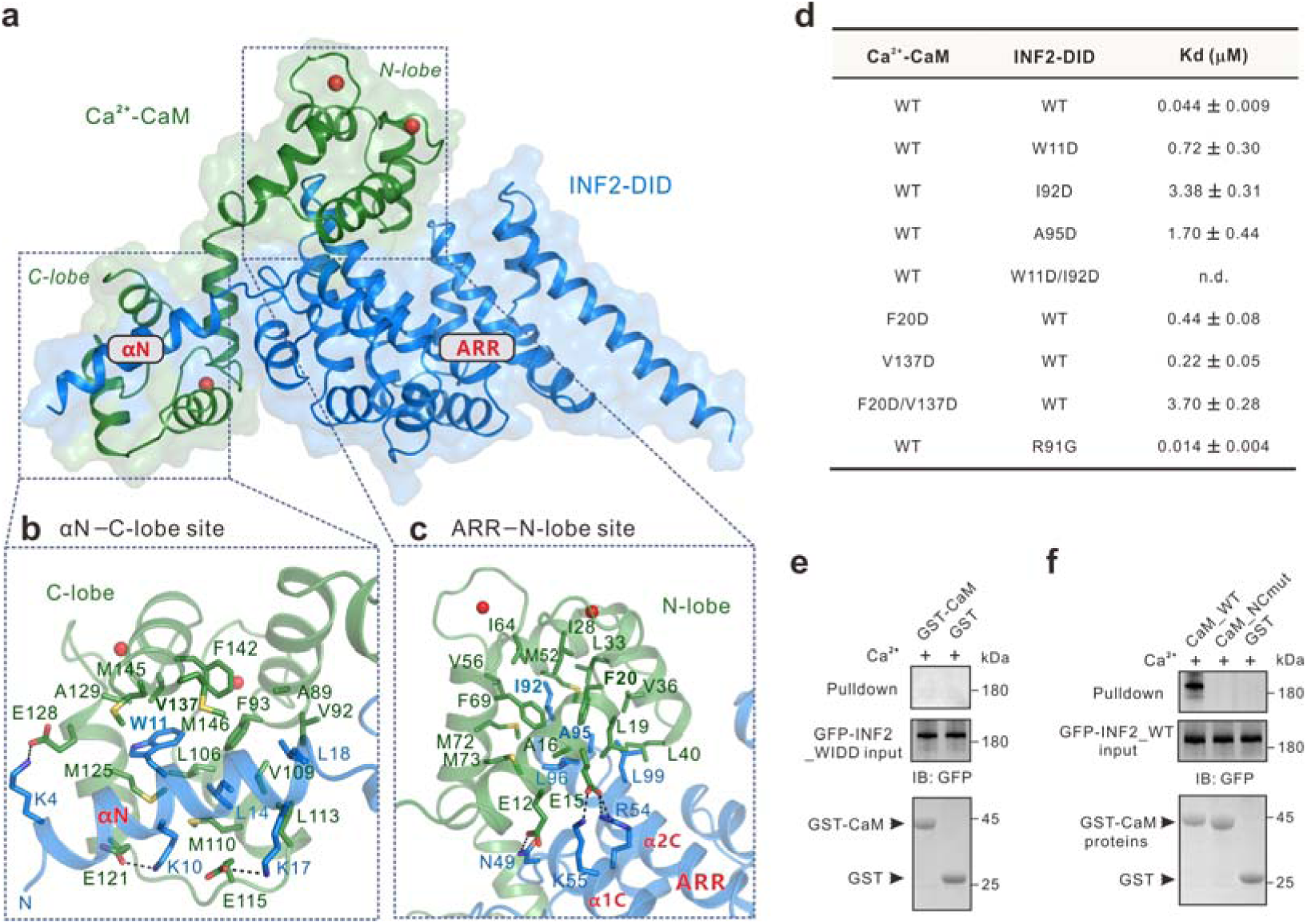
The interaction interface of the INF2 DID–Ca^2+^-CaM complex. **(a)** Combined surface and ribbon representations of the INF2 DID–Ca^2+^-CaM complex structure. The C-lobe–αN and the N-lobe–ARR sites are outlined as dashed boxes. **(b,c)** Detailed views of the C-lobe–αN interface **(b)** and the N-lobe–ARR interface **(c)**. Hydrogen bonds and salt bridges are indicated by dashed lines. **(d)** Summary of ITC-derived binding affinities between WT and mutants of INF2 DID and Ca^2+^-CaM. n.d., not detectable. **(e,f)** GST pull-down assays showing the interactions between CaM and INF2_WIDD **(E)**, and between CaM_NCmut and INF2 **(f)**, in the presence of Ca^2+^. For each pull-down assay, 500□µl of clarified lysate supernatant was incubated for 1□h at 4□°C with 30□µl of GSH-Sepharose 4B slurry beads pre-bound with 2□µM of recombinant GST or GST-CaM proteins, in the presence of 5□mM CaCl_2_.

To evaluate the role of key interface residues in complex formation, we introduced a series of mutations into INF2 DID and CaM. In line with the structural analysis, all mutations significantly impaired the INF2 DID–Ca^2+^-CaM interaction in solution. For example, both the W11D and I92D mutants of INF2 DID reduced binding to Ca^2+^-CaM, while the W11D/I92D double mutant (hereafter referred to as WIDD) completely abolished the interaction, most likely due to disrupted hydrophobic interactions at both sites **(Figs. 2d and S5)**. Reciprocally, the F20D/V137D double mutant of CaM decreased binding affinity to INF2 DID by over 80-fold **(Figs. 2d and S5)**. Further validation showed that GST-tagged Ca^2+^-CaM failed to pull down full-length INF2 bearing the WIDD mutation, confirming the interface observed in the complex structure **(Fig. 2e)**. As a control, the Ca^2+^-binding deficient mutant, CaM_NCmut, also failed to interact with full-length INF2 (Fig. 2f).

### Ca^2+^-CaM binding activates autoinhibited INF2 and promotes actin assembly

We next sought to investigate whether Ca^2+^-CaM directly activates autoinhibited INF2. Like other DRFs, INF2 is autoinhibited through an intramolecular DID–DAD interaction with affinities in the tens of micromolar range ^11,13,21^. Consistently, ITC data confirmed that INF2 DID binds to INF2 DAD with a *K*_d_ value of 34.6 μM **(Fig. 3a)**. This interaction was further confirmed by GST pull-down assays, where GST-tagged INF2 DAD efficiently bound Flag-tagged INF2 DID **(Fig. 3b)**. Crucially, Ca^2+^-CaM disrupted the interaction, whereas the DID_WIDD mutant (deficient in Ca^2+^-CaM binding) retained its binding to DAD even in the presence of Ca^2+^-CaM **(Fig. 3b and 3c)**. These results indicated that binding of Ca^2+^-CaM may disrupt the autoinhibitory DID–DAD interaction.

**Fig. 3.**
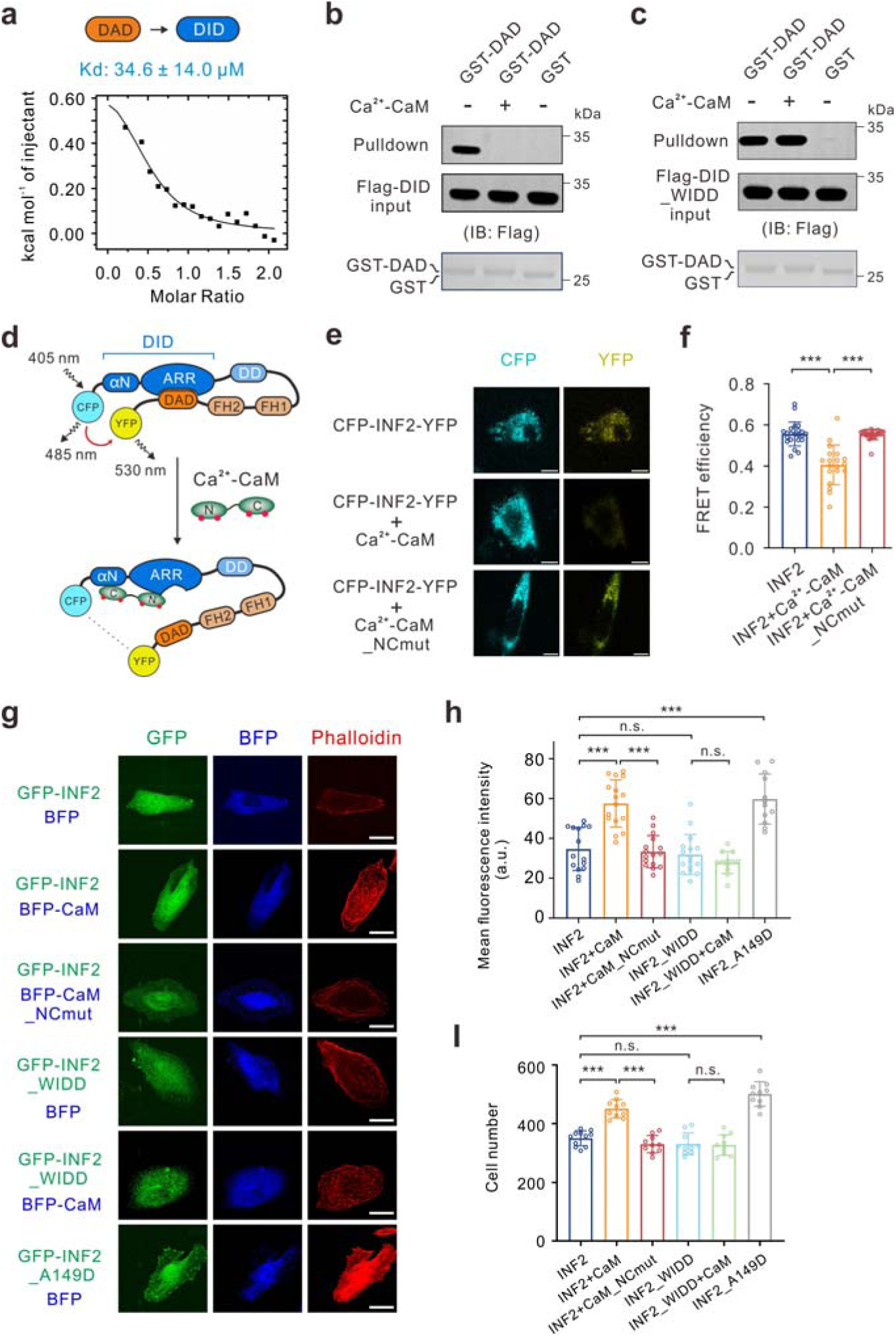
Binding of Ca^2+^-CaM activates INF2 and drives actin assembly. **(a)** ITC-based measurements of the binding affinity between INF2 DID (200 µM) and INF2 DAD (2 mM). **(b,c)** GST pull-down assays showing binding of Ca^2+^-CaM specifically disrupts the DID–DAD interaction **(b)**, but not the DID_WIDD–DAD interaction **(c)**. For each pull-down assay, 500□µl of clarified lysate supernatant was incubated for 1□h at 4□°C with 30□µl of GSH-Sepharose 4B slurry beads pre-bound with 2□µM of recombinant GST or GST-DAD proteins, in the presence or absence of 2□µM Ca^2+^-CaM. **(d)** Schematic of the fluorescence resonance energy transfer (FRET) assay used to monitor intramolecular interactions within INF2 in cells. **(e)** Representative fluorescence images showing that FRET occurs when wild-type HeLa cells was expressed with CFP-INF2-YFP. Co-expression of CaM_WT, but not CaM_NCmut abolished FRET signal. Scale bar, 5 μm. **(f)** Quantification of FRET efficiency in **(e)**. Data were presented as mean ± SD (n = 19 to 22 cells). ***P < 0.001. **(g)** Representative fluorescence images of HeLa cells co-transfected with various GFP-INF2 (non-CAAX) and BFP-CaM constructs. Actin filaments were labeled with phalloidin. Scale bar, 10 μm. **(h)** Quantification of average fluorescence intensity of actin filaments in **(g)**. Data were presented as mean ± SD (n = 12 to 18 cells). ***P < 0.001; n.s., not significant. **(i)** Quantification of number of cells expressing the indicated constructs that have traversed the membrane and are present on its lower surface, counted in each randomly selected microscopic field. Data were presented as mean ± SD from at least ten randomly selected fields from three independent experiments (n = 10). ***P < 0.001; n.s., not significant.

To further validate this mechanism, we employed a fluorescence resonance energy transfer (FRET) assay, a powerful technique widely used to monitor intramolecular interactions ^34^. To this end, we engineered an INF2 FRET sensor by fusing cyan fluorescent protein (CFP) and yellow fluorescent protein (YFP) at the N- and C-termini, respectively **(Fig. 3d)**. In this assay, FRET signals indicate close proximity between the N-and C- termini, consistent with the intramolecular DID–DAD interaction in autoinhibited INF2. Conversely, diminished FRET signals reflect dissociation of the interaction, leading to an extended, active conformation of INF2 **(Fig. 3d)**. Live-cell imaging revealed robust FRET efficiency in cells expressing the INF2 sensor alone, confirming the basal autoinhibited state of INF2 **(Fig. 3e and 3f)**. Strikingly, co-expression of CaM_WT resulted in a significant reduction in FRET efficiency **(Fig. 3e and 3f)**, indicating Ca^2+^-CaM binding disrupts the DID–DAD interaction. This reduction was reversed when cells co-expressed CaM_NCmut, a mutant incapable of binding to INF2 **(Fig. 3e and 3f)**. These data directly demonstrated that Ca^2+^-CaM binding relieves INF2 autoinhibition in physiologically relevant cellular contexts.

We next examined how Ca^2+^-CaM-induced INF2 activation influences actin filament assembly. In HeLa cells, the nonCAAX isoform of INF2 (INF2_WT) exhibited diffuse cytoplasmic distribution and induced actin stress fibers **(Fig. 3g)**, in line with previous studies ^35,36^. The expression levels of all GFP-INF2 constructs used in this assay were confirmed to be comparable by both Western blot analysis and quantitative fluorescence measurement (**Fig. S6a**). Co-expression of CaM_WT, but not CaM_NCmut, significantly enhanced actin stress fiber formation **(Fig. 3g and 3h)**. In contrast, while INF2_WIDD (incapable of binding Ca^2+^-CaM) retained actin assembly activity comparable to INF2_WT, its co-expression with CaM_WT failed to further enhance actin assembly **(Fig. 3g and 3h)**. As a positive control, the constitutively active mutant of INF2 (A149D), which is expected to disrupt the DID–DAD interaction (see below for details), robustly promoted actin assembly even without CaM **(Fig. 3g and 3h)**.

Given the directed actin polymerization is a fundamental driver of membrane protrusion and cell motility, and the established link between INF2 and cancer progression ^37^, we also explored the functional impact of Ca^2+^-CaM-mediated INF2 activation on cell migration. Aligning with its effects on actin assembly, co-expression of CaM_WT with INF2_WT, but not INF2_WIDD, significantly accelerated cell migration **(Figs. 3i and S6b)**. As expected, the constitutively active INF2_A149D drove maximal migration independently of CaM **(Figs. 3i and S6b)**. Collectively, these findings demonstrated that Ca^2+^-CaM binding relieves INF2 autoinhibition, thereby augmenting actin assembly and promoting cell motility.

### Ca^2+^-CaM-induced conformational changes of DID disrupt DID–DAD interaction

How does Ca^2+^-CaM binding activate autoinhibited INF2? Structural comparisons between the INF2 DID–Ca^2+^-CaM complex and the DID–DAD complex may provide mechanistic insights. To this end, we attempted to determine the DID–DAD complex structure. Although crystallization of the complex was unsuccessful, we obtained crystals of apo INF2_DID and solved its structure at 2.39-Å resolution **(Table S1)**. Interestingly, INF2_DID in the asymmetric unit forms a domain-swapped dimer with a symmetry-related molecule in the adjacent asymmetric unit **(Fig. S7a)**. Although full-length INF2 may form a physiological dimer via its canonical dimerization domain (DD, aa 284–363), the INF2_DID construct used for our structural studies lacks the DD and is monomeric in solution **(Fig. S7b)**. Thus, the domain-swapped dimer seen in the asymmetric unit is best interpreted as a non-physiological crystal-packing interface. The overall structure of apo INF2 DID, comprising five armadillo repeats, resembles its conformation in the INF2 DID–Ca^2+^-CaM complex, though the αN helix is not visible in the apo structure **(Fig. S7c)**.

To gain molecular insights into the DID–DAD interaction, we used AlphaFold3 to generate a structural model of the complex **(Fig. S7d)** ^38^. As expected, the model depicts the DAD peptide binding to the concave groove in DID **(Figs. 4a and S7d)**. This predicted, conserved interface provides the necessary reference to contrast with our experimental structure of the Ca^2+^-CaM-bound INF2_DID. Structural superposition of apo DID and the DID–DAD complex revealed that DAD binding does not induce significant conformational changes in DID **(Fig. 4a)** ^5^. In the DID–DAD interface, hydrophobic residues I973, L976, L977, I980, and F984 from the DAD peptide accommodated hydrophobic residues M109, I115, L146, A149, I152, Y153, and L206 in DID. This interface is further reinforced by a hydrogen bond network involving R106^DID^, D974^DAD^, and N110^DID^ **(Fig. 4b)**. It is worth noting that the INF2 DID–DAD complex structure closely resembles that of the mDia1 DID–DAD complex **(Fig. S7e and S7f)** ^5^. Mutagenesis studies validated the interface: DID_A149D, a well-characterized constitutively active variant of INF2, disrupted DID–DAD binding, while DAD_I980D dramatically decreased binding to DID **(Fig. S7g and S7h)**. By contrast, a negative control mutation (DID_I243D), located outside the interface, retained binding to DAD as expected **(Fig. S7g and S7h)**.

**Fig. 4.**
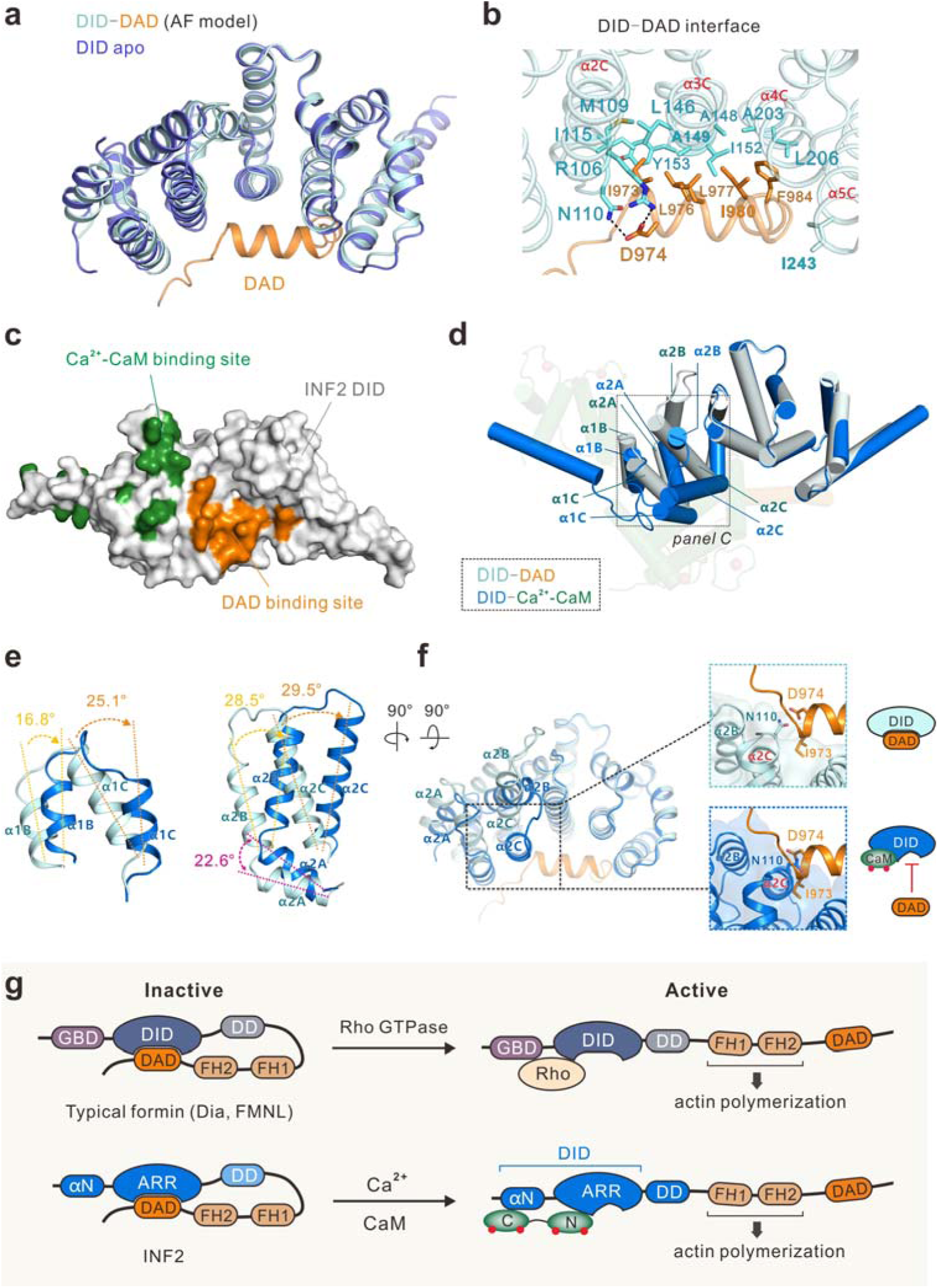
Molecular basis of Ca^2+^-CaM binding-induced allosteric activation of INF2. **(a)** Structural superimposition of INF2 DID (this study) and the DID–DAD complex (predicted by AlphaFold3). **(b)** Detailed interface of the DID–DAD complex. Hydrogen bonds and salt bridges are indicated by dashed lines. **(c)** Surface diagram of binding sites of Ca^2+^-CaM (green) and DAD (orange) on INF2 DID. **(d)** Structural superimposition of the INF2 DID-DAD complex (DID, light cyan; DAD, orange) and INF2 DID–Ca^2+^-CaM complex (DID, blue; Ca^2+^-CaM, green). Helices in INF2 DID that undergo conformational changes upon Ca^2+^-CaM binding are outlined with a dashed box. **(e)** Detailed view of conformational changes in INF2 DID, with angular displacement of each helix indicated. **(f)** Structural superimposition of the DID–DAD and INF2 DID–Ca^2+^-CaM complexes showing Ca^2+^-CaM-induced conformational changes led to a steric clash between the α2C of DID and DAD. **(g)** Depiction of the inhibition and activation mechanisms of formins. Direct binding of Ca^2+^-CaM to INF2 DID induces conformational changes that generate a steric clash between DID and DAD, triggering allosteric activation of INF2. This contrasts with classical formins, which are activated via direct steric displacement of DAD by Rho-GTPase binding.

Having established that these mutations biochemically disrupt the DID–DAD interaction, we next tested whether they would similarly relieve autoinhibition in cells. Indeed, like the constitutively active INF2_A149D, the INF2_I980D mutant drove robust actin assembly and enhanced cell migration **(Fig. S6c-S6f)**. This demonstrates that biochemically disrupting the DID–DAD interface is sufficient to activate INF2’s cellular function.

Since both Ca^2+^-CaM and INF2 DAD bind to INF2 DID, we projected the key binding residues onto the surface of INF2 DID **(Fig. 4c)**. Intriguingly, their interfaces on DID appear largely distinct and non-overlapping **(Figs. 4c and S4)**, suggesting that direct, one-to-one steric competition may not be the primary mechanism. Strikingly, structural superposition of the INF2 DID–Ca^2+^-CaM complex and the INF2 DID–DAD complex revealed that Ca^2+^-CaM binding induces a twisted conformation in INF2 ARR **(Fig. 4d)**. Upon Ca^2+^-CaM binding, N-terminal ARR helices (α1B, α1C, α2A, α2B, and α2C) undergo concerted rotation toward the DAD-binding groove, with angular displacements ranging from 16.8° (α1B) to 29.5° (α2C) **(Fig. 4e)**. These conformational changes led to a steric clash between the α2C helix of DID and the DAD peptide **(Fig. 4f)**. This provides a structural basis for how Ca²⁺-CaM could disrupt autoinhibition, which we biochemically demonstrate by the loss of DID-DAD binding in the presence of Ca^2+^-CaM **(Fig. 3b)**. Taken together, these structural analyses suggested that Ca^2+^-CaM activates autoinhibited INF2 via an allosteric mechanism. This contrasts sharply with Rho-GTPase-mediated activation of classical DRFs, which occurs via direct steric displacement of DAD **(Fig. 4g)** ^7,8^. It remains an open question whether other formins are also regulated by this calcium-dependent allosteric activation mechanism.

### Ca^2+^-CaM-induced activation of INF2 promotes mitochondrial fission

A key step in mitochondrial fission is the activation of ER-bound INF2, which initiates actin polymerization at ER-mitochondrial contact sites ^28^. While ER-released calcium transients have been implicated in this process ^27,39,40^, how they activate INF2 during mitochondrial fission remains unclear. Building on our structural and biochemical evidence that Ca^2+^-CaM directly relieves INF2 autoinhibition, we tested whether CaM acts as a Ca^2+^ sensor to activate INF2 during mitochondrial fission.

Consistent with previous studies, expression of the constitutively active INF2_A149D-CAAX variant (INF2_A149D) in INF2-deficient cells significantly facilitated mitochondrial fission, compared with INF2_WT (**Fig. 5a and 5b**) ^15^. Notably, co-expression of CaM_WT (but not CaM_NCmut) with INF2_WT fully recapitulated this effect, reducing mitochondrial length and increasing fission **(Fig. 5a and 5b)**. By contrast, while INF2_WIDD cells exhibited mitochondrial length comparable to INF2_WT, co-expression of CaM_WT failed to induce mitochondria shortening **(Fig. 5a and 5b)**, indicating that intact Ca^2+^-CaM binding is essential for INF2 activation in this context. These findings aligned precisely with our actin assembly data **(Fig. 3)**, providing compelling evidence that Ca^2+^-CaM-induced INF2 activation drives mitochondrial fission via actin cytoskeleton remodeling.

**Fig. 5.**
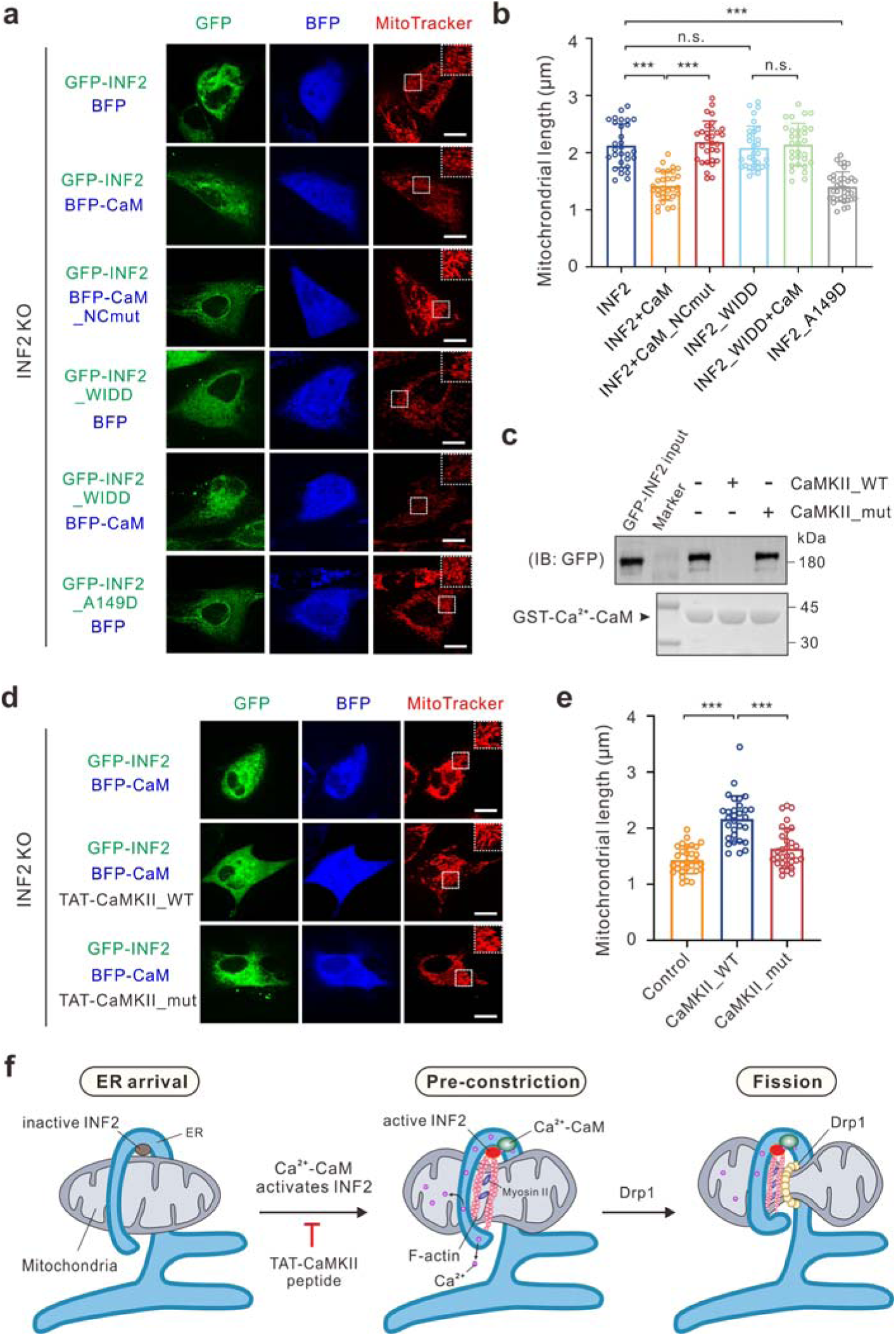
Ca^2+^-CaM-induced activation of INF2 promotes mitochondrial fission. **(a)** Representative fluorescence images of INF2 KO HeLa cells rescued with indicated GFP-INF2 (CAAX) and BFP-CaM constructs. Mitochondria were labeled with MitoTracker. Scale bar, 10 μm. **(b)** Quantification of average mitochondrial length in **(a)**. For each condition, at least 40 mitochondria from each of 30 cells (n = 30 cells) were traced end-to-end. All measurements were conducted by an observer blinded to the experimental conditions. Data were presented as mean ± SD. ***P < 0.001; n.s., not significant. **(c)** GST pull-down assay showing that the CaMKII_WT peptide, but not the CaMKII_mut peptide disrupted the INF2–Ca^2+^-CaM interaction. For each pull-down assay, 500□µL of clarified lysate supernatant was incubated for 1□h at 4□°C with 30□µL of GSH-Sepharose 4B slurry beads pre-bound with 2□µM GST-Ca^2+^-CaM, in the presence of 2□µM CaMKII_WT peptide or CaMKII_mut peptide. **(d)** Representative fluorescence images of INF2 KO HeLa cells rescued with WT GFP-INF2 and BFP-CaM, treated with TAT-CaMKⅡ_WT (10 μM) or TAT-CaMKⅡ_mut peptides (10 μM). Mitochondria were labeled with MitoTracker. Scale bar, 10 μm. **(e)** Quantification of average mitochondrial length in **(d)**. Data were presented as mean ± SD (n = 30 cells). ***P < 0.001; n.s., not significant. **(f)** Schematic model of INF2’s function during mitochondrial fission. Fission is initiated when ER wraps around mitochondria, inducing pre-constriction at their contacts. Mitochondrial pre-constriction is triggered by calcium-mediated allosteric activation of ER-bound INF2, which drives actin polymerization around ER. These actin filaments recruit non-muscle myosin II to constrict mitochondria, followed by Drp1 recruitment to complete fission.

To further validate the CaM–INF2 axis in mitochondrial fission, we developed a competitive inhibitory peptide strategy. Based on the conserved CaM-binding motif of calcium/calmodulin-stimulated protein kinase II (CaMKII) **(Fig. S8a)** ^41^, we designed wild-type (CaMKII_WT) and mutant (CaMKII_mut) peptides. GST pull-down assays confirmed Ca^2+^-dependent CaM binding to CaMKII_WT but not CaMKII_mut **(Fig. S8b and S8c)**. Crucially, CaMKII_WT (but not CaMKII_mut) competitively disrupted the Ca^2+^-CaM–INF2 interaction **(Fig. 5c)**, confirming this peptide as a specific molecular tool to probe INF2 regulation.

To evaluate the impact of these peptides on mitochondrial fission, we conjugated the CaMKII_WT and CaMKII_mut peptides with the cell-penetrating human immunodeficiency virus transactivator of transcription (TAT) peptide. The peptides were designed to be identical except for the two functional residues **(Fig. S8b)**, ensuring comparable physicochemical properties for cellular entry. In INF2-deficient cells co-expressing INF2_WT and CaM, treatment of TAT-CaMKII_WT, but not TAT-CaMKII_mut, significantly attenuated mitochondrial fission, yielding elongated mitochondria **(Fig. 5d and 5e)**. Together, these results established CaM as the essential Ca^2+^ sensor that activates INF2 to drive mitochondrial fission **(Fig. 5f)**.

### CMT-associated INF2 mutation confers gain-of-function by enhanced Ca^2+^-CaM binding

Over 70 pathogenic mutations of INF2 have been identified in patients with FSGS and CMT **(Fig. 6a and Fig. S9a)** ^42^. Interestingly, all known pathogenic INF2 missense mutations cluster within exons encoding the DID ^42^. The INF2 DID–Ca^2+^-CaM complex structure and the Ca^2+^-dependent allosteric activation mechanism uncovered here allow us to interpret the molecular basis of these disease mutations. Mapping these variants onto the complex structure revealed that most are localized to the folding core of INF2 DID **(Fig. 6a)**, potentially disrupting its integrity.

**Fig. 6.**
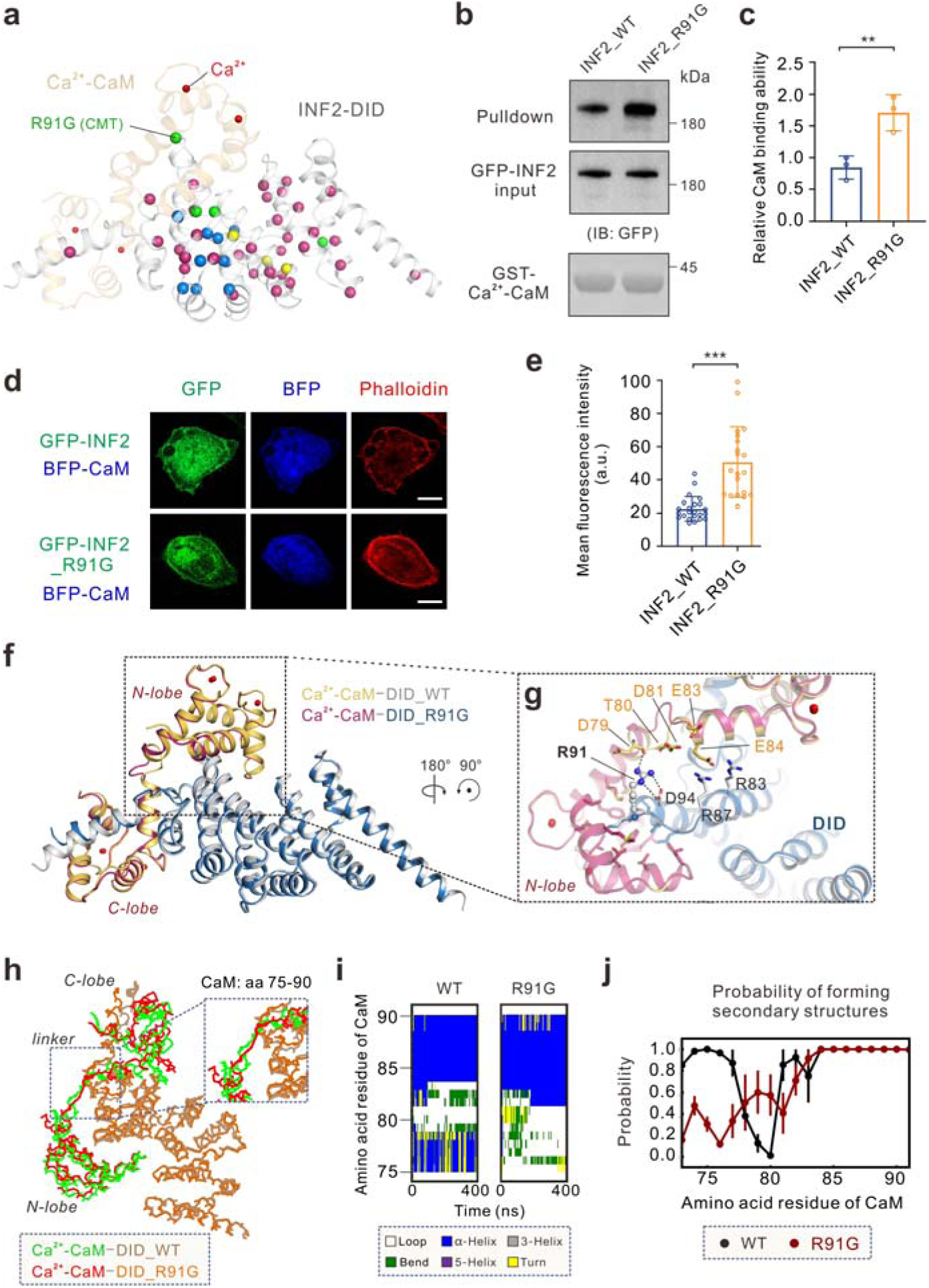
A CMT-associated INF2 mutation confers gain-of-function effect by enhanced Ca^2+^-CaM binding. **(a)** Mapping of disease-associated mutations onto the INF2 DID–Ca^2+^-CaM complex structure. **(b)** GST pull-down assay showing enhanced Ca^2+^-CaM binding by INF2_R91G compared to INF2_WT. **(c)** Quantification of relative Ca^2+^-CaM binding ability in **(b)**. Data were presented as mean ± SD. **P <0.01. **(d)** Representative fluorescence images of HeLa cells co-transfected with indicated GFP-INF2 (nonCAAX) and BFP-CaM constructs. Actin filaments were stained by phalloidin. Scale bar, 10 μm. **(e)** Quantification of average fluorescence intensity of actin filaments in **(d)**. Data were presented as mean ± SD (n = 20 to 23 cells). ***P < 0.001. **(f)** Structural superimposition of the DID_WT–Ca^2+^-CaM (CaM, gray; DID_WT, yellow) and the DID_R91G–Ca^2+^-CaM complex (CaM, red; INF2-DID, blue) structures. **(g)** Structural comparison showing that key residues at the N-lobe–ARR interface is largely conserved between the two complexes. Of note, R91 forms a salt bridge with INF2_D94 and a hydrogen bond with the mainchain carbonyl of CaM_D79. This local network helps stabilize the positioning of CaM’s acidic cluster (D81, E83, E84), enabling their optimal interaction with INF2 basic residues (R83, R87). These interactions promote a more rigid and stable α-helical conformation in CaM_linker (aa 75-90). **(h)** Comparison of the average structures of the DID_WT–Ca^2+^-CaM complex (CaM, green; DID_WT, brown) and the DID_R91G–Ca^2+^-CaM complex (CaM, red; INF2-DID, orange) from molecular dynamics (MD) simulations. Noted that the linker of CaM (aa 75-90) adopts different conformations in two complex structures. **(i)** The secondary structure of Ca^2+^-CaM as a function of simulation time. A representative result of multiple replicates is shown. **(j)** Probability of secondary structure formation in Ca^2+^-CaM (aa 73–91) when bound to DID_WT versus DID_R91G during MD simulations. Data represent the average of three parallel simulations, with error bars indicating the standard error of the mean.

The CMT-associated R91G mutation of INF2 attracts our attention because of its proximity to the N-lobe–ARR interface **(Fig. 6a)** ^43^. We set to assess the impact of this variant on the INF2–Ca^2+^-CaM interaction. Unexpectedly, DID_R91G binds to Ca^2+^-CaM with a *K*_d_ value of 14 nM, nearly threefold stronger than DID_WT **(Fig. 1c *vs* Fig. s9b)**. GST pull-down assays confirmed the enhanced complex formation **(Fig. 6b and 6c)**, indicating R91G thermodynamically stabilizes the INF2–CaM interaction. Consistent with hyperactivation, INF2_R91G exhibited markedly elevated actin polymerization upon CaM co-expression **(Fig. 6d and 6e)**. These data established R91G as a pathological gain-of-function mutation that potentiates INF2 activation via enhanced Ca^2+^-CaM binding.

How does INF2_R91G enhance Ca^2+^-CaM binding? To address this question, we determined the crystal structure of the DID_R91G–Ca^2+^-CaM complex at 2.11-Å resolution **(Fig. S9c, S9d and Table S1)**. Surprisingly, structural superposition of the DID_WT–Ca^2+^-CaM complex and the DID_R91G–Ca^2+^-CaM complex revealed near-identical conformations (root-mean-square deviation, RMSD = 0.298 Å; **Fig. 6f)**. Key interface interactions are largely conserved between the two complexes **(Fig. 6g)**. The only difference lies at position 91: in the WT complex, the side chain of R91 forms a local salt bridge with INF2_D94 and a hydrogen bond with the mainchain carbonyl of CaM_D79 **(Fig. 6g)**.

This discrepancy implies that the enhanced affinity stems from altered conformational dynamics rather than static structural changes. We hypothesized that the R91G substitution could optimize the dynamic ensemble of the INF2–CaM complex by modulating interfacial dynamics. To test this, we employed molecular dynamics (MD) simulations to characterize conformational dynamics of both complexes. MD simulations confirmed that the R91G complex exhibits greater overall stability than the wild-type complex, with CaM in the WT complex deviating more readily from its initial conformation **(Fig. S9e)**. This enhanced stability most likely arises from altered dynamics of the inter-lobe linker of CaM (aa 75-90, CaM_linker). In the R91G complex, the CaM_linker sampled a broader range of conformations with a higher propensity for dynamic loops, whereas it was more restrained in the wild-type complex **(Figs. 6h-6j and S9f).** Quantitative analysis revealed a significantly elevated RMSD for the R91G complex, confirming the enhanced flexibility of the CaM_linker **(Fig. S9g)**. Thus, by destabilizing the rigidifying local network, the R91G mutation promotes tighter interfacial packing and greater global complex stability via a dynamics-driven mechanism.

## Discussion

Formins regulate actin cytoskeleton remodeling, enabling eukaryotic cells to alter their shape and behavior in response to diverse signals. Calcium serves as a critical link between these signals and actin dynamics. For decades, however, our understanding of calcium-mediated actin polymerization has focused on indirect signaling cascades: calcium-activated enzymes (kinases or phosphatases) that converge on regulators of Rho GTPases, which in turn modulate formin-mediated actin assembly ^44^. A critical unresolved question remains whether spatial calcium signals can bypass these multi-step cascades to directly instruct actin assembly.

Here, we resolve this question by providing the first atomic-resolution mechanistic blueprint for direct calcium-to-actin signal transduction. We demonstrate that Ca^2+^-CaM directly binds to INF2 DID with high affinity, inducing allosteric conformational changes that sterically disrupt INF2 autoinhibition, thereby activating INF2 to facilitate actin assembly. This direct activation is enabled by a unique bipartite binding interface where Ca^2+^-CaM simultaneously engages INF2’s αN helix (via its C-lobe) and ARR (via its N-lobe). Cooperative saturation of CaM with four Ca^2+^ ions (two per lobe) generates a nonlinear activation threshold, which filters out basal calcium fluctuations and underpins the system’s ultrasensitive response to localized calcium spikes within confined subcellular compartments.

This mechanism achieves precise spatiotemporal decoding of calcium microdomains across the cell. At ER-mitochondria contact sites, it permits activated ER-tethered INF2 to trigger burst-like actin assembly exclusively at sites of ER calcium release, thereby driving mitochondrial fission **(Fig. 5f).** Notably, a recent study have shown that ER-tethered INF2 also executes endosome/lysosome fission via actin polymerization, establishing this formin as a universal calcium–actin transducer at organelle contact interfaces ^45^. Therefore, this Ca^2+^-CaM–mediated allosteric activation of INF2 represents an evolutionarily optimized regulatory design: stringent intrinsic autoinhibition prevents ectopic activation, while cooperative Ca^2+^ sensing ensures rapid responses to organelle remodeling demands.

Notably, INF2’s regulation may extend beyond this direct Ca^2+^-CaM-mediated pathway, involving additional layers of control that may modulate its activity in specific cellular contexts. Recent studies have shown that in certain contexts, a complex of cyclase-associated protein (CAP) and lysine-acetylated actin (KAc-actin) is required to reinforce DID–DAD-mediated autoinhibition ^46^. HDAC6-mediated actin deacetylation relieves this “facilitated autoinhibition”, contributing to INF2 activation ^46,47^. The interplay between intrinsic and facilitated autoinhibition constitutes a hierarchical regulatory mechanism that fine-tunes INF2 sensitivity across distinct subcellular microdomains. For example, at ER microdomains, ER-bound INF2 encounters minimal CAP/KAc-actin stabilization (CAP is cytosolic), rendering it hyperresponsive to ER-localized calcium bursts via Ca^2+^-CaM-mediated activation mechanism. In the perinuclear cytosol, by contrast, elevated CAP/KAc-actin levels may raise the INF2 activation threshold, probably requiring coincident calcium signals and HDAC6 activity, a ‘two-key’ safeguard against spurious activation from global calcium waves. Future work should explore how calcium signaling and posttranslational modifications cooperate within this integrated regulatory framework.

Dysregulation of this direct activation pathway might be a central driver of INF2-associated human disease. Most pathogenic INF2 mutations localize to its DID domain and are thought to destabilize the domain’s structural integrity, leading to aberrant actin cytoskeletal organization. In contrast to the loss-of-function mutations, the gain-of-function effects of the CMT-associated INF2 R91G mutation are particularly striking. It enhances Ca^2+^-CaM binding and constitutively activates actin polymerization. Intriguingly, this increased Ca^2+^-CaM binding does not stem from changes in static structural features but rather from optimized dynamic interactions at the INF2 DID_R91G–Ca^2+^-CaM interface, highlighting the critical role of conformational dynamics in protein complex assembly. It is plausible that hyperactivated INF2 R91G could induce abnormal cell motility and/or mitochondrial dysfunction, contributing to the onset and progression of CMT. This hypothesis requires direct experimental testing in the future. This gain-of-function model aligns with observations for other INF2 mutations linked to FSGS (e.g., S186P and R218Q), which also induce INF2 hyperactivation, drive aberrant actin organization, and trigger excessive mitochondrial fission ^48^. Notably, our CaMKII-derived inhibitory peptide, which disrupts the Ca^2+^-CaM–INF2 interaction, could directly counter these pathological effects by normalizing hyperactive INF2, thus offering a precision strategy for INF2-linked diseases like CMT and FSGS.

In summary, our work defines the Ca^2+^-CaM–INF2 axis as a direct molecular transducer that converts spatial calcium signals into localized actin assembly. These findings advance our understanding of how cells encode spatial calcium information into actin-dependent processes, and establishes the INF2–CaM axis as a novel therapeutic target for modulating aberrant actin cytoskeletal dynamics in various diseases, including neurodegenerative disorders and renal diseases.

## Acknowledgments

We thank beamlines BL02U1, BL18U1 and BL19U1 at Shanghai Synchrotron Radiation Facility (SSRF, China) for X-ray beam time, the staff members of the Large-scale Protein Preparation System, Nuclear Magnetic Resonance System, and Molecular Imaging System at the National Facility for Protein Science in Shanghai, for providing technical support and assistance in data collection and analysis. We also thank Mingjie Zhang for valuable discussion and critical comments on the manuscript. This work was supported by the STI2030 Major Projects (2022ZD0214400), the National Natural Science Foundation of China (32470728) as well as the Natural Science Foundation of Chongqing, China (CSTB2023NSCQ-JQX0031) to J.Zhu, the ‘Science and Technology Innovation Action Plan’ of the Shanghai Science and Technology Commission (24ZR1433500) to L.L, the Natural Science Foundation of Jiangsu, China (BK20240784) and the Natural Science Foundation for Higher Education Institutions of Jiangsu, China (24KJB180018) to M.Z.

## Author contributions

B.Z., M.Z., L.L., C.F., and J.Zhu conceptualized and designed the experiments. B.Z. M.Z. and L.L. performed biochemical assays and structural studies. K.L., C.Z. and C.F. conducted mitochondrial imaging experiments and analyses. J.Zhang and R.G. carried out molecular dynamic simulations and related analyses. Z.L and Y.F. provided technical support and assistance. All authors contributed to data analysis and interpretation. J.Zhu drafted the manuscript with input from all authors and supervised the project.

## Data availability

The atomic coordinates of apo INF2 DID, the INF2 DID_WT–Ca^2+^-CaM complex and the INF2 DID_R91G–Ca^2+^-CaM complex have been deposited to the Protein Data Bank under the accession codes 9W3X, 9W46, and 9W4G, respectively. All other data are available from the corresponding author upon reasonable request.

## Methods

### Constructs and peptides

The coding sequences of various fragments of INF2 (NM_001374199.1) and Calmodulin (CaM, NM_017326.3) were cloned into a modified pET32a vector containing an N-terminal Trx-His₆ tag. For GST pull-down assays, various fragments of INF2 and CaM were cloned into the pGEX-4T-1 vector. For fluorescence resonance energy transfer (FRET) assays, full-length INF2 was cloned into a pPBbsr2-Raichu vector to generated a CFP-INF2-YFP FRET sensor. All mutations were generated by standard PCR-based mutagenesis method using the Phanta Max super fidelity DNA polymerase (Vazyme, P505). All constructs were confirmed by DNA sequencing.

The wild-type TAT-CaMKII peptide (YGRKKRRQRRRARRKLKGAILTTMLATRNF) and TAT-CaMKII_mut peptide (YGRKKRRQRRRARRKDKGADLTTMLATRNF) were commercially synthesized by ChinaPeptides (Shanghai, China) with purity > 95%.

### Protein Expression and Purification

Recombinant proteins were expressed in *Escherichia coli* BL21 (CodonPlus) or BL21 (DE3) cells at 16□°C for 18 h, induced with 0.2 mM isopropyl-β-D-thiogalactoside (IPTG). The N-terminal Trx-His6 tagged and GST-tagged proteins were purified by Ni^2+^-NTA agarose affinity chromatography and GSH-Sepharose affinity chromatography (Cytiva), respectively. Proteins were further purified by size-exclusion chromatography (SEC, Cytiva) in buffer containing 50 mM Tris pH 8.0, 100 mM NaCl, 5 mM CaCl_2_, and 4 mM β-Mercaptoethanol (β-ME). Trx-tagged proteins were cleaved by the human rhinovirus 3C protease at 4 °C overnight and the Trx-His₆ tag was then removed by an additional SEC step.

### Crystallization, Data collection and Structure determination

All crystals were obtained by the sitting-drop vapor diffusion method at 16 °C, with each drop composed of 0.5 μl of protein complex (∼20 mg/ml) and 0.5 μl of reservoir solution. The crystals of the INF2 DID_WT–Ca^2+^-CaM and INF2 DID_R91G–Ca^2+^-CaM complexes were grown in the buffer containing 100 mM MES pH 6.5, 200 mM Ammonium sulfate, 30% PEG5000 MME. The crystal of apo INF2 DID was grown in 0.05 M Ammonium sulfate, 0.05 M BIS-TRIS pH 6.5, 30% v/v Pentaerythritol ethoxylate (15/4 EO/OH). Crystals were soaked in crystallization solution containing 20% glycerol for cryoprotection.

Diffraction data were collected at Shanghai Synchrotron Radiation Facility (SSRF, Shanghai, China) and were processed with XDS ^49^. All crystal structures were solved via the molecular replacement method implemented in PHASER ^50^, with specific searching models as follows: the apo INF2 DID structure was solved using the mDia1 DID structure (PDB code: 2BNX) as the search model; the INF2 DID_WT–Ca^2+^–CaM complex structure was solved by employing both the apo INF2 DID structure and the Ca^2+^–CaM structure (PDB code: 3GP2) as search models; the INF2 DID_R91G–Ca^2+^–CaM complex structure was solved using the INF2 DID_WT–Ca^2+^–CaM complex structure as the search template. Subsequent structural refinement was performed iteratively using PHENIX ^50^ refinement and Coot ^51^. Final refinement statistics of all crystal structures were provided in Table S1. Structural diagrams were generated using PyMOL.

### GST pull-down assay

GFP- or Flag-tagged INF2 fragments were overexpressed in HEK293T cells. Cells were harvested and lysed on ice using lysis buffer (10□mM HEPES pH 7.5, 100□mM NaCl, 0.1□mM MgCl_2_, 1% Triton X-100, supplemented with a protease inhibitor cocktail; 1 mL per 10 cm dish). Lysates were clarified by centrifugation at 14,000□g for 10□min at 4□°C. For each pull-down, 500□µL of clarified supernatant was incubated for 1□h at 4□°C with 30□µL of GSH-Sepharose 4B slurry beads pre-bound with 2□µM of recombinant GST, wild-type GST-CaM, or the indicated GST-CaM mutants, in the presence or absence of 5□mM CaCl_2_. Following incubation, beads were collected and washed twice with 500 µl of ice-cold lysis buffer. Bound proteins were eluted by boiling the beads in 30□μL 2× SDS-PAGE loading dye. Eluted proteins were resolved by SDS-PAGE and analyzed by western blot using an anti-GFP (Santa Cruz, 1:5,000) and anti-Flag (Proteintech; 1:2,000) antibodies.

### Isothermal titration calorimetry (ITC)

ITC measurements were performed using a MicroCal iTC200 system (Malvern Panalytical, UK) at 25 °C. All proteins were dissolved in titration buffer containing 50 mM Tris pH 8.0, 100 mM NaCl, 5 mM CaCl_2_, and 4 mM β-ME. The syringe was loaded with the titrant protein (∼500 μM), and 2 μl aliquots were injected into the sample cell containing the target protein (∼50 μM) at 120 s intervals to allow complete equilibration between injections. Raw data were analyzed using the Origin 7.0 software package (Microcal).

### Analytical gel filtration chromatography coupled with static light scattering

Size-exclusion chromatography with multi-angle light scattering analysis was performed at 25 °C on an Agilent InfinityLab HPLC system coupled with a miniDawn static light scattering detector (Wyatt) and an Optilab differential refractive index detector (Wyatt). Protein samples (100 μM in 300 μL) were loaded into a Superose 12 10/300 GL column (Cytiva) pre-equilibrated with running buffer (50 mM Tris pH 8.0, 100 mM NaCl, 5 mM CaCl_2_, and 4 mM β-ME). Data collection and analysis were performed using ASTRA 6 (Wyatt).

For analytical gel filtration chromatography used to characterize the interaction between Ca^2+^-CaM and INF2_DID, the assay was carried out on an AKTA FPLC system (GE Healthcare) at 25 °C. Protein samples (50 μM in 300 μL) were loaded into a Superdex 200 10/300 GL column (Cytiva) pre-equilibrated with running buffer.

### Cell culture

HEK293T (ATCC) and HeLa (ATCC) cell lines were cultured in Dulbecco’s Modified Eagle Medium (DMEM, Gibco) containing 10% fetal bovine serum (FBS, Gibco), supplemented with 100 U/ml penicillin, and 100 μg/ml streptomycin (Gibco) at 37□°C and 5% CO_2_.

### Generation of knockout cell lines

*INF2* knockout HeLa cells were generated using the CRISPR-Cas9 system. Briefly, a plasmid encoding the Cas9 protein and a single guide RNA (sgRNA) targeting exon 2 of the human *INF2* gene (5’-CGGAGATACGTGCAACGCCG-3’) was transfected into HeLa cells via electroporation. Following transfection, cells were subjected to monoclonal isolation. Genomic DNA was extracted from monoclonal colonies, and the targeted region was amplified by PCR. The amplified PCR products were then verified by Sanger sequencing.

### Fluorescence resonance energy transfer (FRET)

HeLa cells were co-transfected with 0.5□μg of the CFP-INF2-YFP construct with Flag-tagged CaM (WT or mutant) using Lipofectamine 2000 (Invitrogen). After 24 h post-transfection, FRET measurements were conducted using a Zeiss LSM 900 confocal laser scanning microscope. For CFP fluorescence visualization, transfected cells were excited at 405nm, and emission signals were collected at 485 nm (CFP) and 530 nm (YFP). FRET efficiency (*E*) was calculated using the sensitized emission method with background correction, calculated as: *E*=*I*_YFP_/(*I*_CFP_+*I*_YFP_), where *I*_YFP_ and *I*_CFP_ are the background-subtracted fluorescence intensities of the YFP and CFP channels, respectively.

### Immunofluorescence staining for actin filaments

HeLa cells grown on 12 well plates were transfected with 0.5□μg of indicated plasmids using Lipofectamine 2000 (Invitrogen). After 24 h post-transfection, cells were fixed in 4% (w/v) formaldehyde freshly prepared from paraformaldehyde in PBS for 10 min at room temperature, followed by permeabilization with 0.5% (v/v) Triton-X 100 in PBS for 5 min. For actin filament staining, cells were incubated with TRITC-conjugated Phalloidin (Yeasen) for 25 min at room temperature. The immunofluorescence images were acquired using a Leica TCS SP8 confocal microscope with consistent laser power and gain settings across samples. Quantitative analysis of actin filament levels was performed by measuring the average fluorescence intensity of phalloidin staining using ImageJ software, with background fluorescence subtracted from each image prior to analysis. All measurements were conducted by an observer blinded to the experimental conditions. Data were presented as mean ± SD from three independent experiments.

### Transwell migration assay

Cell migration was assessed using Corning Costar transwell membrane filter inserts (8-μm pore size). HeLa cells transfected with indicated plasmids were serum-starved for 4 h, then resuspended in serum-free DMEM and seeded into the upper chamber (1 ∼ 1.5×10□ cells per well). The lower chamber contained DMEM supplemented with 10% FBS as a chemoattractant. Following incubation at 37□°C for 16 to 18 h, non-migrated cells on the upper surface of the membrane were gently removed with a cotton swab. Migrated cells on the lower surface were fixed with 100% methanol for 15 min, washed with PBS, and then stained with 0.1% Crystal Violet Staining Solution (Sangon Biotech) for 15 min. The cell migration images were acquired using a 10 × light microscope. Data were presented as mean ± SD by counting the number of migrated cells in at least 10 randomly selected fields from three independent experiments.

### Mitochondrial staining and Live-cell imaging

At 48 h post-transfection with 1□μg of indicated plasmids via Lipofectamine 3000 (Invitrogen), HeLa cells grown on glass-bottom dishes were incubated with 100 nM MitoTracker™ Red CMXRos (Thermo Fisher Scientific) in pre-warmed phenol-red-free DMEM for 15 min at 37 °C in the dark. Live-cell imaging was performed using a PerkinElmer UltraVIEW spinning disk confocal microscope equipped with an environmental chamber set to maintain physiological temperature and CO_2_. All images were acquired with a 0.8 μm step size containing 11 planes. Laser power and detector gain were kept consistent across all samples. Mitochondrial length was measured using the freehand line tool in ImageJ processing program. At least 40 mitochondria per cell (30 cells per condition) were measured to calculate the average length. Flat cellular regions with clearly resolvable mitochondria were selected. For each condition, at least 40 mitochondria from each of 30 cells were traced end-to-end. All measurements were conducted by an observer blinded to the experimental conditions. For the TAT-peptide treatment, 10□μM of either the TAT-CaMKII_WT or TAT-CaMKII_mut peptide dissolved in PBS was added to the cells that had been transfected with indicated plasmids. Mitochondrial length was measured following the same method as described above. Data were presented as mean ± SD from three independent experiments.

### Molecular dynamics simulations

Molecular dynamics simulations of Ca^2+^-CaM in complex with either WT or R91G mutant of INF2 DID were performed using GROMACS 2022.2 package ^52^ with the CHARMM36m force field ^53^. The force field parameters of Ca^2+^ ions were taken from the multi-site ion model ^54^. Systems were solvated in cubic boxes with TIP3P water, with counter ions added to neutralize the simulation systems. The systems were subjected to energy minimization using the steepest descent algorithm, which was followed by 1 ns equilibrium simulations with the backbone constrained with a force constant of 1000 kJ/mol/nm^2^. Production runs consisted of three independent 400 ns simulations per system, using a 2 fs integration time step. Bonds involving hydrogen atoms were constrained with the LINCS algorithm ^55^; long-range electrostatics were calculated via the Particle Mesh Ewald method ^56^, with a Fourier mesh grid spacing of 0.12 nm, an interpolation order of 4 and a cutoff value of 1.2 nm, and Van der Waals interactions used a force-switch cutoff (1.0-1.2 nm). The V-rescale thermostat ^57^ was used to maintain the temperature of the system at 300 K, with a relaxation time of 1 ps, while the isotropic C-rescale barostat ^58^ was employed to restrain the pressure of the system at 1 bar, using a compressibility of 4.5×10^-5^ and a relaxation time of 1 ps.

### Statistical analysis

Statistical parameters are reported in the Figures and corresponding Figure Legends. For transwell migration assays, immunofluorescence staining of actin filaments, mitochondrial staining and FRET assays, the results were expressed as mean ± SD; ns, not significant, ∗∗∗p < 0.001, ∗∗p < 0.01, using one-way analysis of variance (ANOVA) test. For GST pull-down assays, the results were expressed as mean ± SD; ns, not significant, ∗∗∗p < 0.001, ∗∗p < 0.01, using two-tailed student’s *t* test. Statistical analysis was performed by GraphPad Prism. All experiments were performed at least three times.

