## Supplementary material for "Calcium directs actin assembly via allosteric activation of formin INF2": Figures S1 to S9 and Tables S1

**This PDF file includes:**

Figures S1 to S9

Tables S1

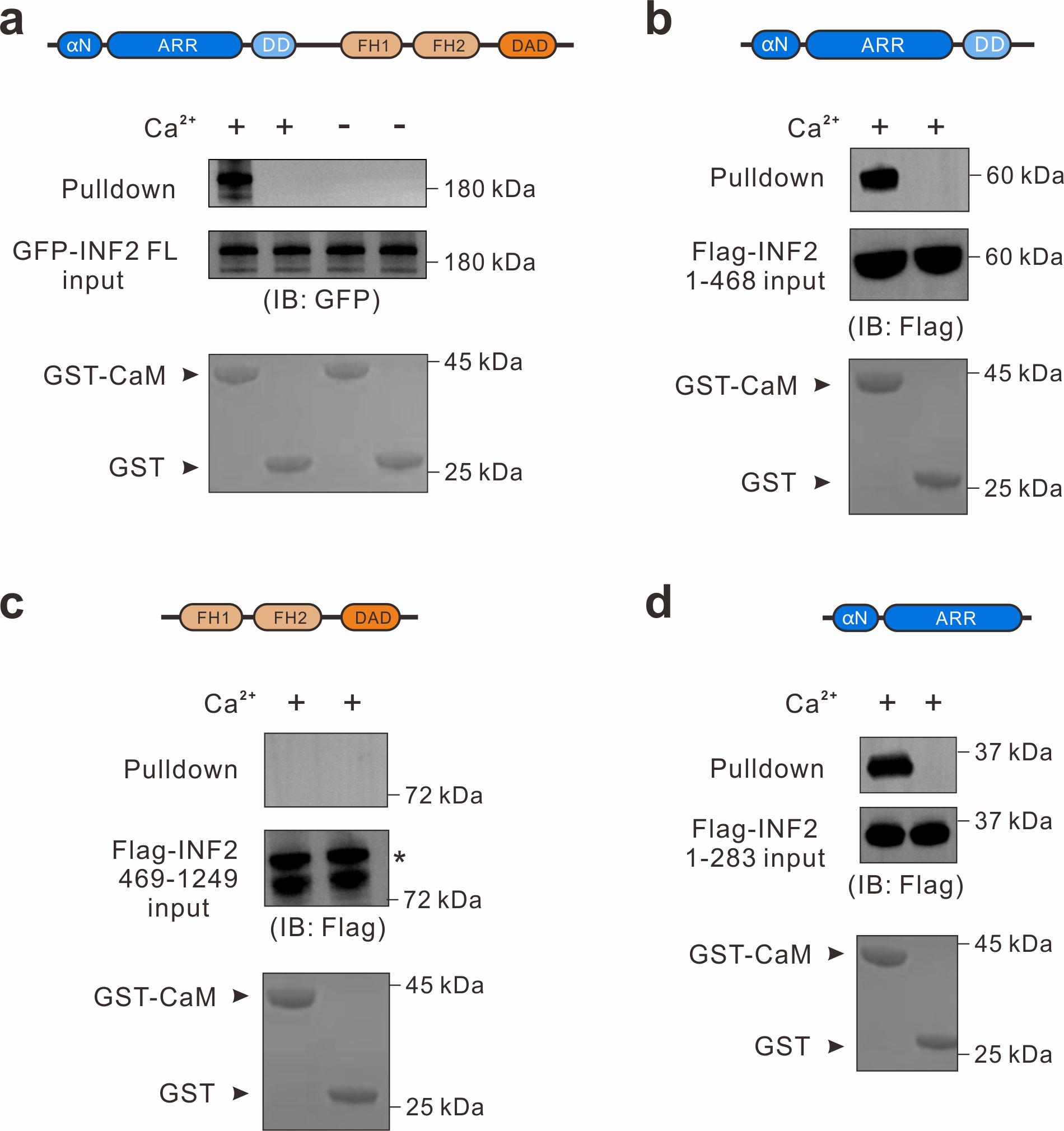

**Fig. S1. Characterization of the binding between INF2 and CaM.**

**(a)** GST pull-down assay showed that full-length INF2 binds to CaM in the presence of Ca^2+^. **(b-d)** GST pull-down assays showed that Ca^2+^-CaM specifically binds to INF2 DID-DD (aa 1-468) **(b)** and INF2 DID (aa 1-283) **(d)**, but not INF2 FH1-FH2-DAD tandem (aa 469-1249) **(c)**. For each pull-down assay, 500 µl cell lysate supernatant was incubated for 1 h at 4 °C with 30 µl of GSH-Sepharose 4B slurry beads pre-bound with 2 µM GST-CaM or GST, in the presence or absence of 5 mM CaCl_2_. Following incubation, beads were collected and washed twice with 500 µl of ice-cold lysis buffer. Bound proteins were eluted and then resolved by SDS-PAGE and analyzed by western blot.

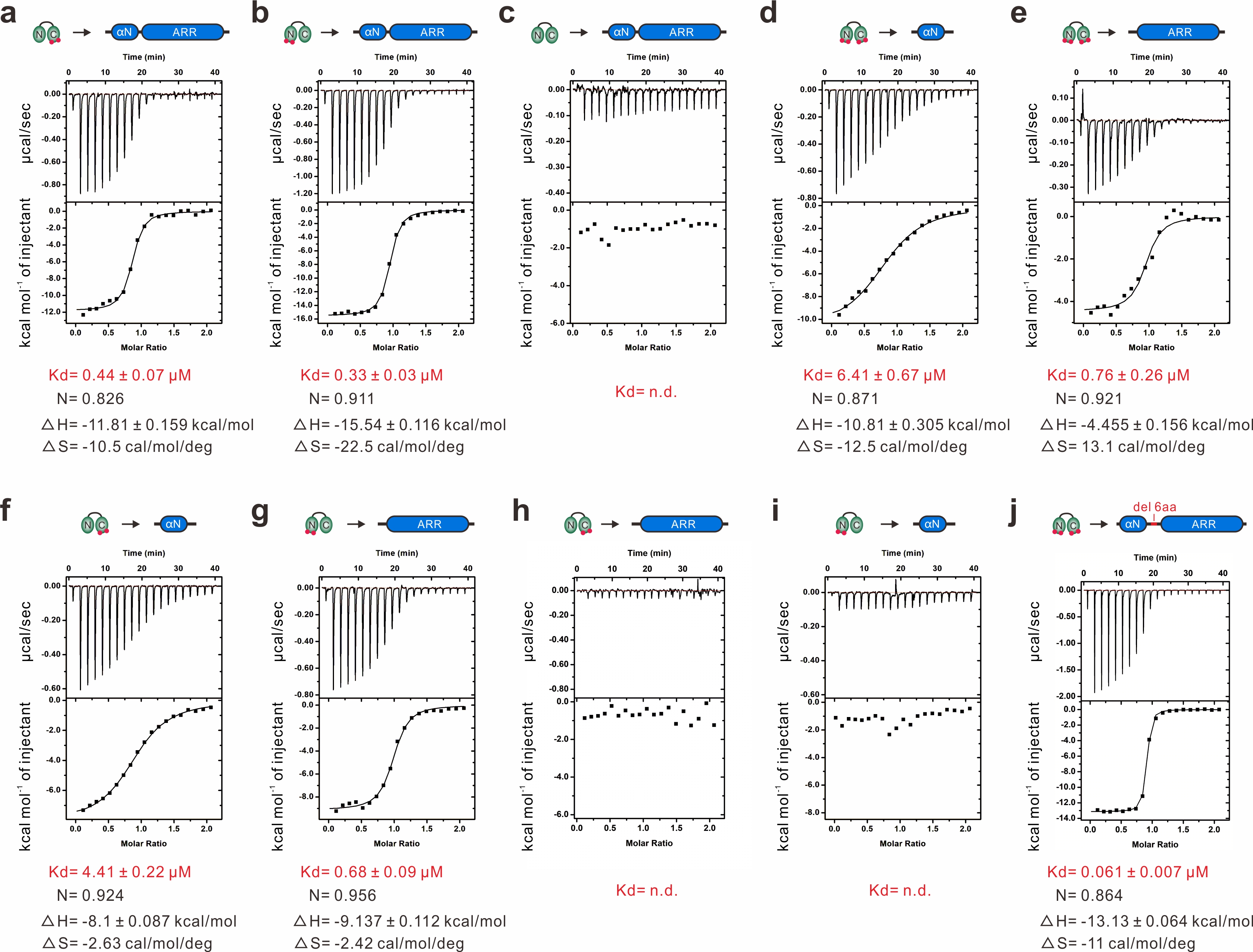

**Fig. S2. ITC-based characterization of the interactions between INF2 and CaM in the presence of Ca^2+^.**

ITC-based measurements of the binding affinities between CaM N-lobe_mut and INF2 DID **(a)**, CaM C-lobe_mut and INF2 DID **(b)**, CaM_NCmut and INF2 DID **(c)**, CaM and INF2 αN **(d)**, CaM and INF2 ARR **(e)**, CaM N-lobe_mut and INF2 αN **(f)**, CaM C-lobe_mut and INF2 ARR **(g)**, CaM N-lobe_mut and INF2 ARR **(h)**, CaM C-lobe_mut and INF2 αN **(i)**, CaM and INF2 DID^del6^ **(j)**. Noted that the concentration of CaM (wild-type or mutants) and various INF2 fragments were 500 μM and 50 μM, respectively. n.d., not detectable.

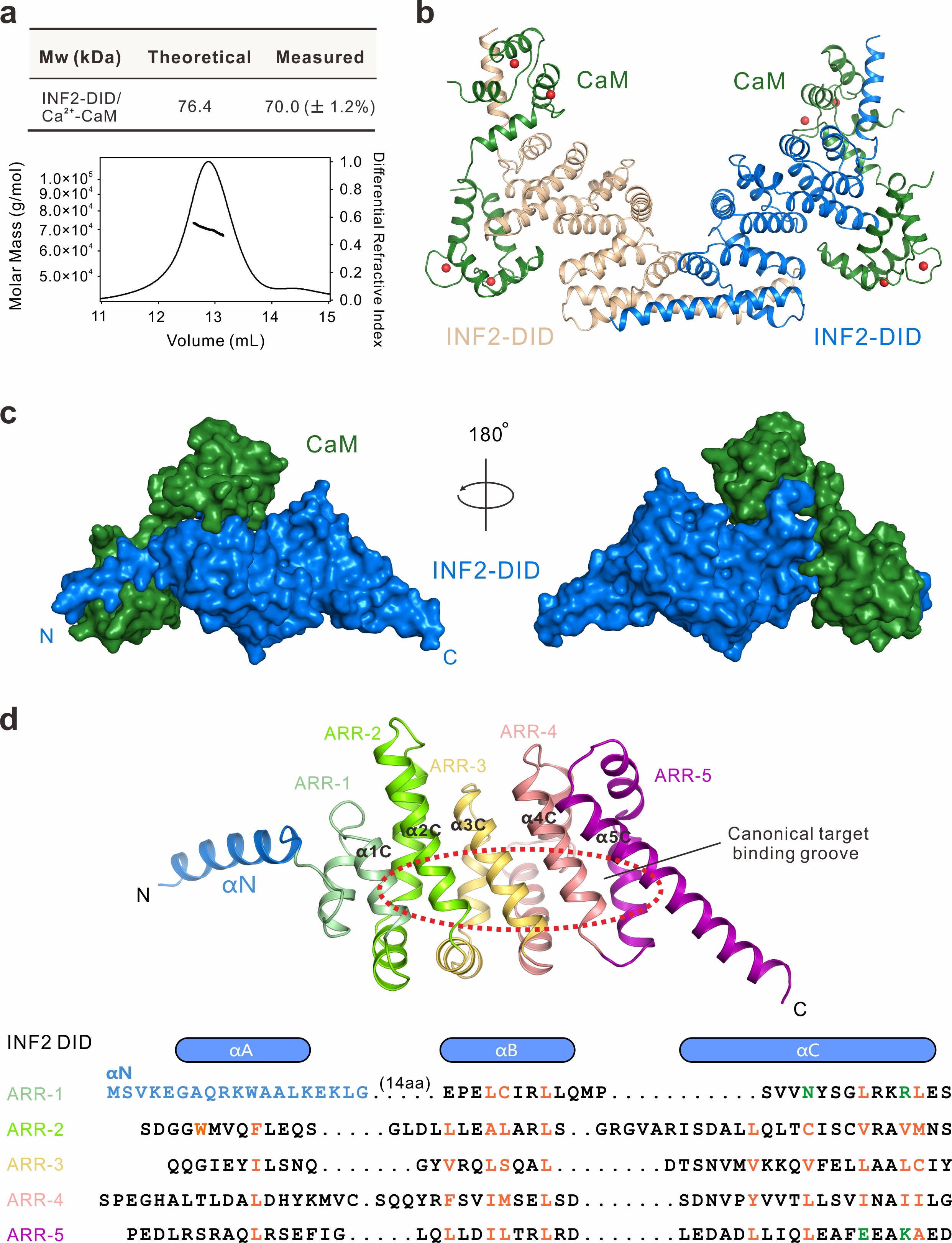

**Fig. S3. Structural analysis of the INF2 DID–Ca^2+^-CaM complex.**

**(a)** Analytical gel filtration chromatography coupled with static light scattering assay showing a 1:1 stoichiometry of the INF2 DID–Ca^2+^-CaM complex (100 μM). **(b)** Each asymmetric unit of the crystal contains two INF2 DID–Ca^2+^-CaM complex molecules. **(c)** Surface representation shows the overall architecture of the INF2 DID–Ca^2+^-CaM complex. **(d)** Overall structure and structure-based sequence alignment of armadillo repeats in INF2. Hydrophobic residues that mediate the core scaffold within the armadillo repeats are marked orange. Polar residues in the αC of first and fifth repeat are labeled green. αN of INF2 DID is labeled blue. Canonical target binding groove of ARR is shown by a red dashed circle.

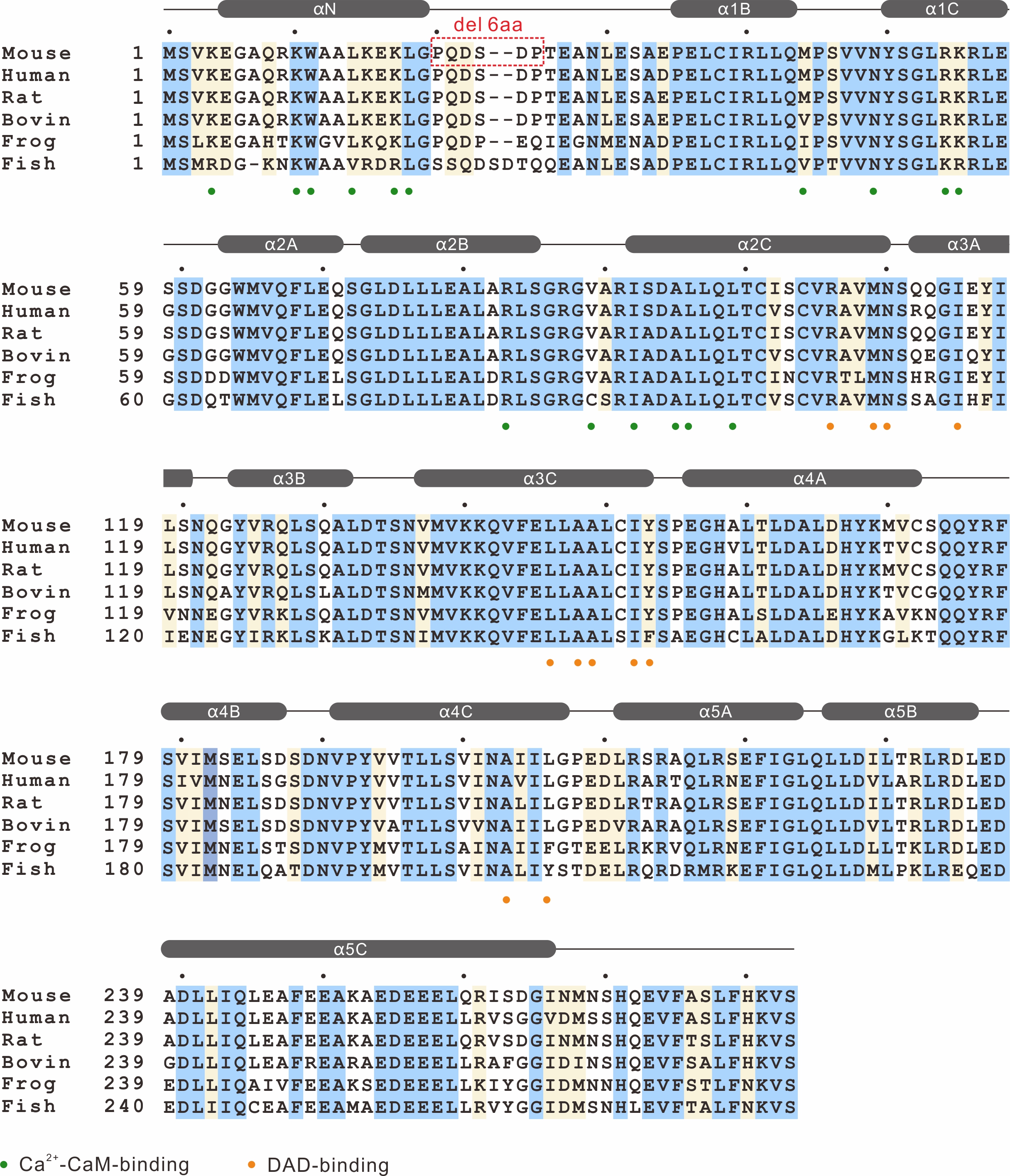

**Fig. S4. Sequence analysis of INF2 DID.**

Amino acid sequence alignment of INF2 DID from different species. In this alignment, absolutely conserved amino acids are highlighted in blue, and highly conserved residues are highlighted in yellow. Residues involved in the binding to Ca^2+^-CaM are indicated with green dots. Residues involved in the INF2 DAD binding are indicated with orange dots. Secondary structures of INF2 DID are indicated at the top.

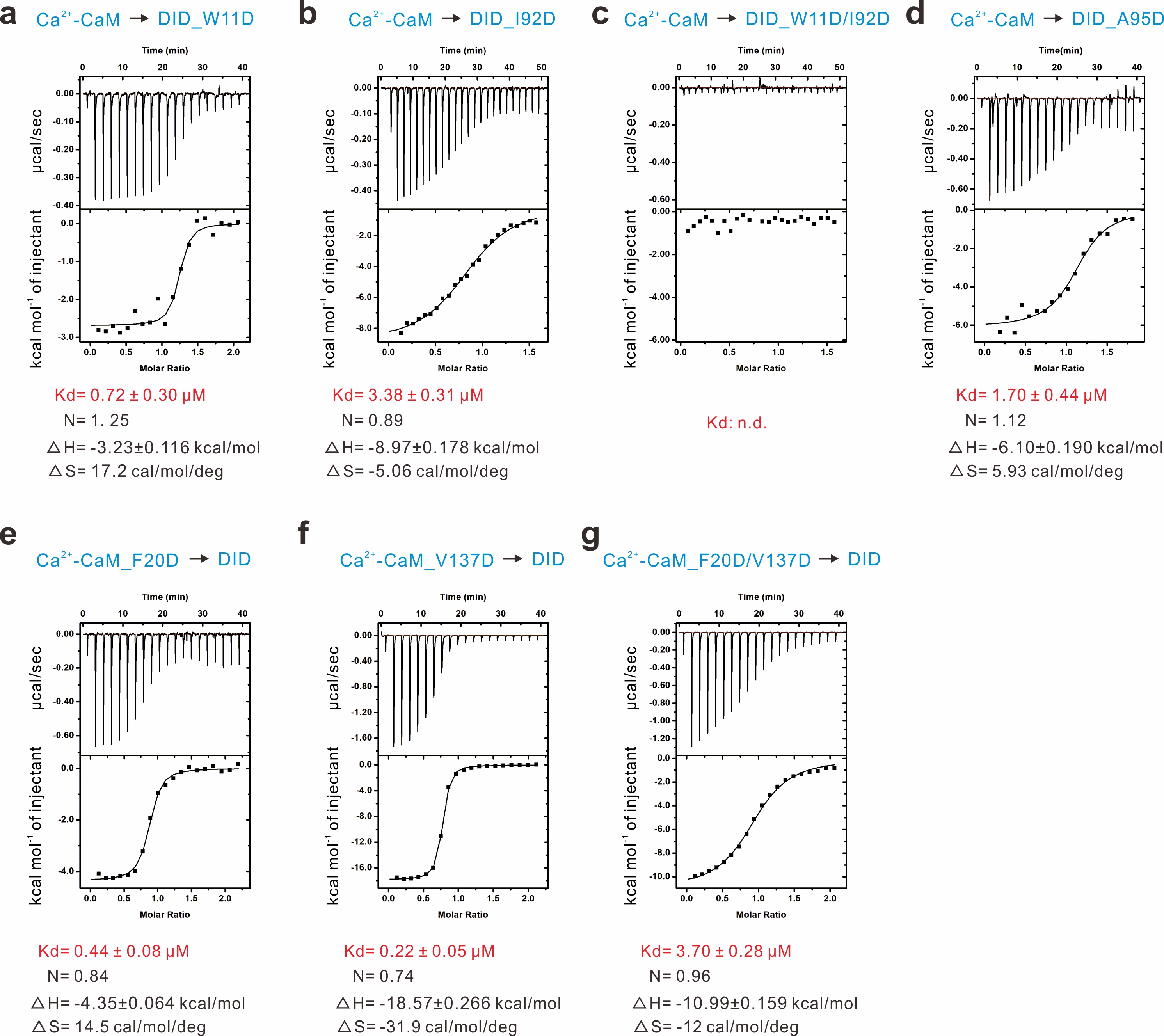

**Fig. S5. ITC-based characterization of various mutations of INF2 and Ca^2+^-CaM.**

ITC-based measurements of the binding affinities between Ca^2+^-CaM and DID_W11D **(a)**, Ca^2+^-CaM and DID_I92D **(b)**, Ca^2+^-CaM and DID_WIDD **(c)**, Ca^2+^-CaM and DID_A95D **(d)**, Ca^2+^-CaM_F20D and DID **(e)**, Ca^2+^-CaM_V137D and DID **(f)**, Ca^2+^-CaM_ F20D/V137D and DID **(g)**. Noted that the concentration of CaM (wild-type or mutants) and INF2 DID (wild-type or mutants) were 500 μM and 50 μM, respectively. n.d., not detectable.

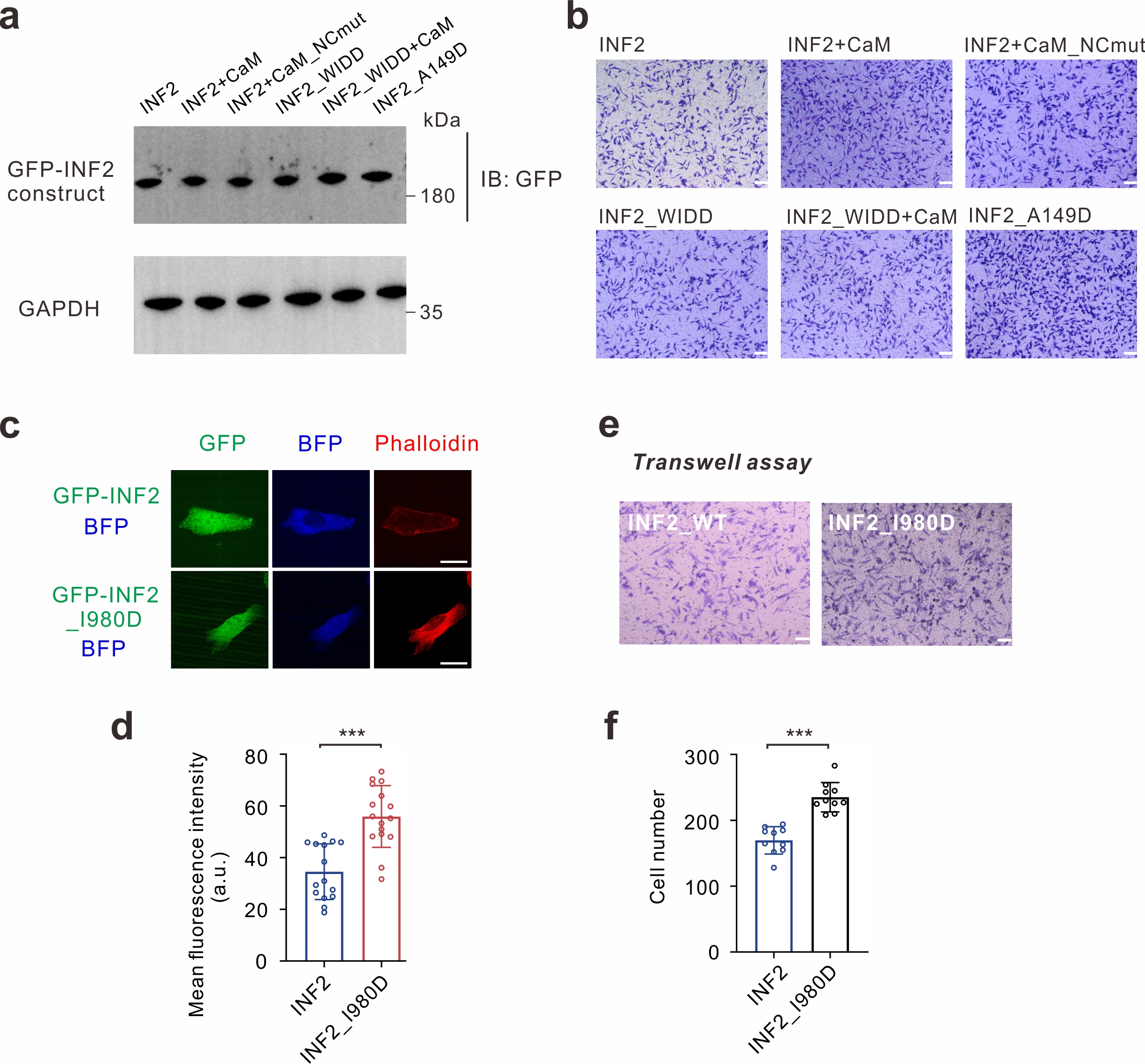

**Fig. S6. Functional characterization of the INF2 DID–Ca^2+^-CaM complex and autoinhibited INF2.**

**(a)** Western blot analysis verified that the different GFP-INF2 constructs were expressed at comparable levels. **(b)** Representative cell migration images of HeLa cells overexpressed with different GFP-INF2 and BFP-tagged CaM plasmids. Scale bar, 50 μm. **(c)** Representative images showing replacement of I980 by Asp in INF2 significantly promoted actin assembly in HeLa cells. Actin filaments were labeled with Phalloidin. Scale bar, 10 μm. **(d)** Quantification of average fluorescence intensity of actin filaments in in HeLa cells expressed the indicated constructs. Data were presented as mean ± SD (n = 15 cells). ***P <0.001. **(e)** Transwell migration assays were performed to measure the cell migration activities of HeLa cells transfected with indicated constructs. Scale bar, 50 μm. **(f)** Quantification of number of cells expressing the indicated constructs that have traversed the membrane and are present on its lower surface, counted in each randomly selected microscopic field. Data were presented as mean ± SD from at least ten randomly selected fields from three independent experiments (n = 10). ***P <0.01.

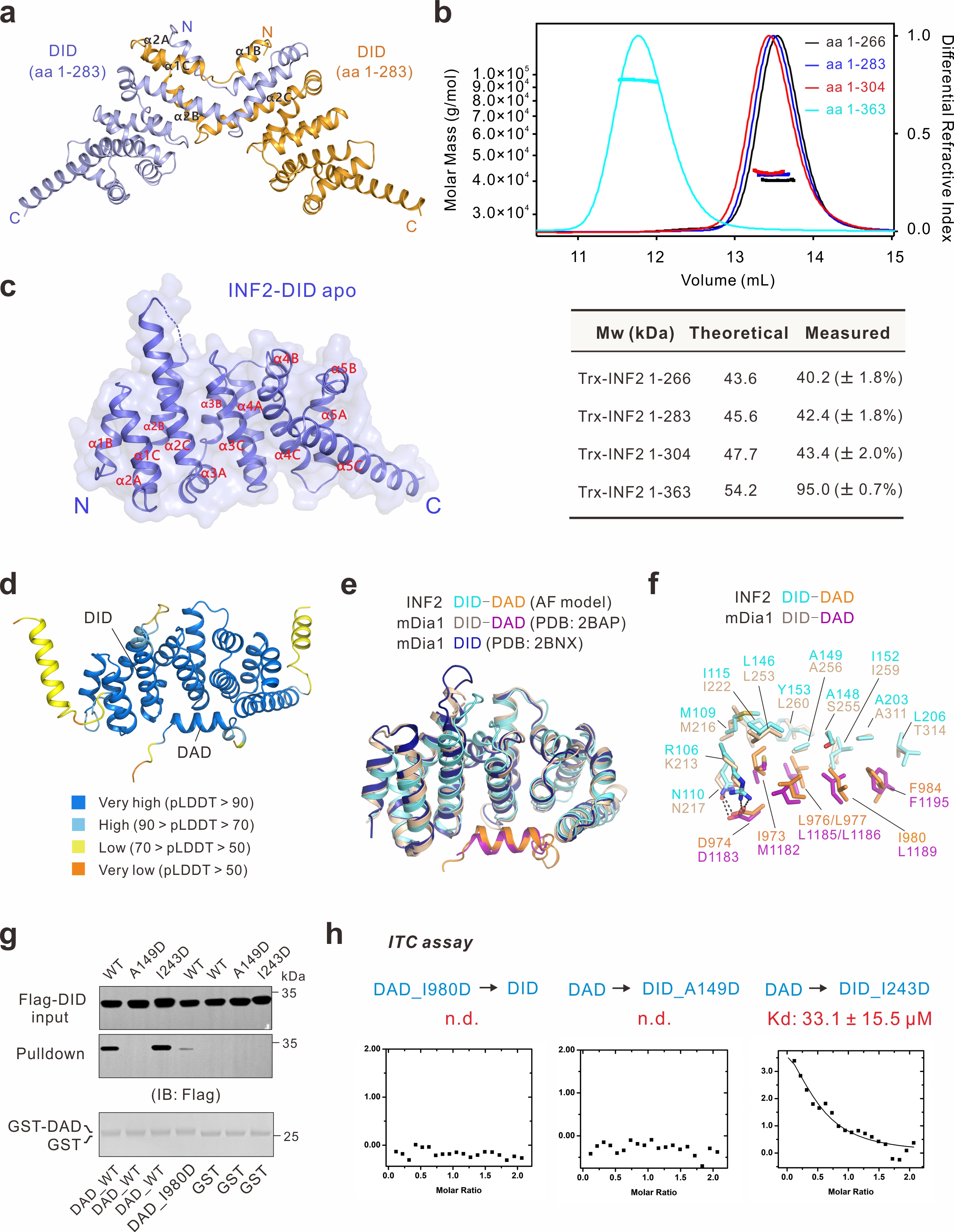

**Fig. S7. Structural and functional analysis of the INF2 DID–DAD interaction.**

**(a)** Ribbon diagram of the overall structure showing that INF2 DID (aa 1-283) exists as a swapped dimer in the crystal structure. **(b)** Analytical gel filtration chromatography coupled with static light scattering assay showing various fragments of INF2 (100 μM) with different molecular weights in solution. **(c)** Combined surface and ribbon representations of the crystal structure of apo INF2 DID. **(d)** Ribbon diagram of the overall structure of the INF2 DID–DAD complex predicted by AlphaFold3. **(e)** Structural superimposition of INF2 DID–DAD complex (AlphaFold3 model), mDia1 DID–DAD complex (PDB code: 2BAP) and mDia1 DID apo structure (PDB code: 2BNX). **(f)** Superimposition of the key residues for DID–DAD interactions in INF2 and mDia1. **(g)** GST pull-down assays showing that key residues involved in the DID–DAD interface are required for the intact interaction. For each assay, 500 µl cell lysate supernatant was incubated for 1 h at 4 °C with 30 µl of GSH-Sepharose 4B slurry beads pre-bound with 2 µM GST-DAD or GST. Following incubation, beads were collected and washed twice with 500 µl of ice-cold lysis buffer. Bound proteins were eluted and then resolved by SDS-PAGE and analyzed by western blot. **(h)** ITC-based measurements of the binding affinities between DAD_I980D and DID, DAD and DID_A149D, DAD and DID_I243D. Noted that the concentration of DAD (wild-type or mutants) and DID (wild-type or mutants) were 2 mM and 200 μM, respectively. n.d., not detectable.

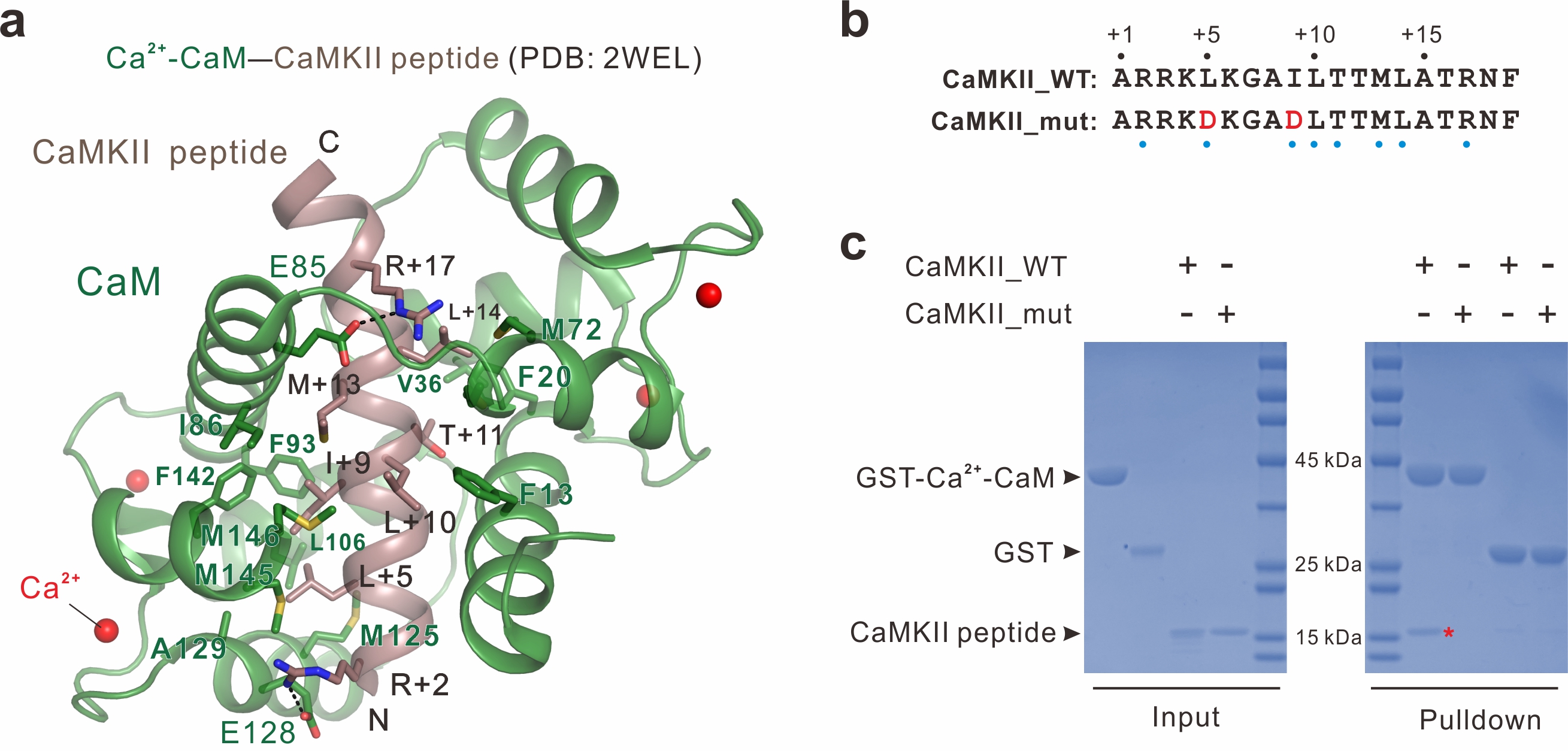

**Fig. S8. Design of a CaMKII inhibitory peptide capable of disrupting the INF2–Ca^2+^-CaM complex.**

**(a)** Detailed interface of the Ca^2+^-CaM–CaMKII peptide complex (PDB code: 2WEL). Dotted lines denote hydrogen bonds and salt bridge interactions. **(b)** Amino acid sequence alignment of CaMKII_WT and CaMKII_mut. The residues involved in the binding to Ca^2+^-CaM were indicated with blue dots. **(c)** GST pull-down assays showing that Ca^2+^-CaM (2 μM) binds to trx-tagged CaMKII_WT (2 μM), but not trx-tagged CaMKII_mut (2 μM).

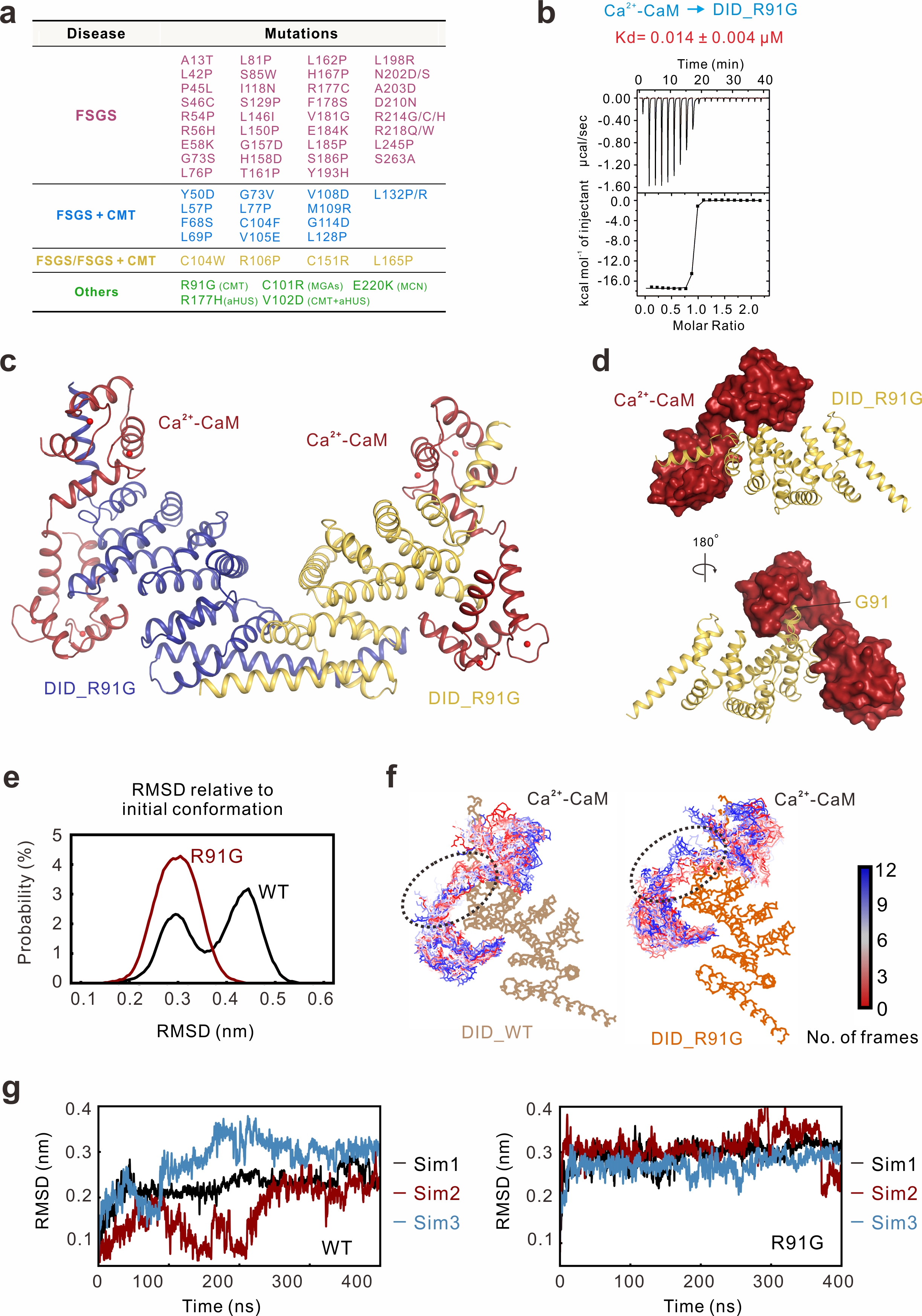

**Fig. S9. Structural analysis of the INF2-DID_R91G–Ca^2+^-CaM complex.**

**(a)** Summary of INF2-associated disease mutations: isolated FSGS (red), FSGS (in some) or FSGS/CMT (in others; yellow), FSGS/CMT (blue), and other diseases (green). **(b)** ITC analysis of bindings of INF2 DID_R91G (50 μM) to Ca^2+^-CaM (500 μM). **(c)** Ribbon diagram of the overall structure showing that INF2-DID_R91G–Ca^2+^-CaM complex exists as a dimer in the crystal structure. **(d)** Combined surface and ribbon representations of the INF2-DID_R91G–Ca^2+^-CaM complex. **(e)** Distributions of the root-mean-square deviation (RMSD) values in molecular dynamics (MD) simulations of Ca^2+^-CaM in complex with either INF2 DID_WT or INF2 DID_ R91G mutant. The initial backbone structure of each complex served as the reference frame for RMSD calculation. **(f)** Representative simulation snapshots taken at 100 ns intervals (total 12 frames) were superimposed and aligned based on the backbone atoms of INF2_DID to illustrate conformational sampling. Ca^2+^-CaM is shown in a color gradient from blue (early frames) to red (late frames). Visual comparison reveals that the central linker region of Ca^2+^-CaM, when bound to the R91G DID mutant, explores a broader range of conformations compared to its complex with the WT DID, indicating higher conformational variability. **(g)** To quantify linker flexibility, the RMSD of the central Ca^2+^-CaM linker was calculated over time for each complex (three simulations for each complex). The results establish that the CaM_linker exhibits a consistently elevated RMSD in the R91G mutant complex compared to the WT complex, confirming a mutation-induced enhancement of conformational dynamics.

| **Table S1. Data collection and refinement statistics** | | | |
| --- | --- | --- | --- |
| **Data collection and processing** | | | |
| **Crystal** | **INF2 DID** | **INF2 DID_WT–Ca^2+^-CaM** | **INF2 DID_R91G–Ca^2+^-CaM** |
| Source | SSRF-BL18U1 | SSRF-BL19U1 | SSRF-BL19U1 |
| Wavelength(Å) | 0.96183 | 0.97851 | 0.97853 |
| Space group | R32 | P2_1_ | P2_1_ |
| Unit cell (a,b,c,Å) | 84.35, 84.35, 242.66 | 37.19, 110.92, 113.58 | 36.99, 110.59, 112.66 |
| Unit cell (α, β, γ, °) | 90, 90, 120 | 90, 91.88, 90 | 90, 91.87, 90 |
| Resolution range (Å) | 29.19-2.39 (2.45-2.39) | 79.35-2.11(2.22-2.11) | 30.84-2.11(2.16-2.11) |
| No. of unique reflections | 13552 (979) | 53164 (7721) | 51665 (3839) |
| Redundancy | 18.5 (15.9) | 6.5 (6.5) | 3.4 (3.5) |
| I/σ(I) | 13.6 (1.7) | 10.1 (2.5) | 11.2 (1.8) |
| Completeness (%) | 99.9 (99.6) | 99.4 (99.2) | 99.1 (98.7) |
| R_merge_ (%)^a^ | 13.3 (185.6) | 10.2 (79.8) | 8.6 (86.4) |
| CC1/2 | 99.8 (70.8) | 99.7 (79.3) | 99.8 (56.1) |
| **Structure refinement** | | | |
| R_work_^b^/R_free_^c^ (%) | 22.54/26.86 | 19.32/24.14 | 19.67/22.79 |
| rmsd bonds (Å)/angles (°) | 0.009/1.160 | 0.010/1.150 | 0.008/1.010 |
| Number of reflections | | | |
| Working set | 12756 | 52789 | 51649 |
| Test set | 1277 | 2570 | 2682 |
| Number of protein atoms | 1854 | 6466 | 6389 |
| Average B factor protein/solvent (Å^2^) | 66.36/56.83 | 49.64/48.73 | 41.69/44.09 |
| Clashscore | 3.22 | 4.75 | 4.18 |
| Ramachandran plot(%) | | | |
| Most favored regions | 98.30 | 99.38 | 99.14 |
| Additionally allowed | 1.70 | 0.62 | 0.86 |
| Outliers | 0 | 0 | 0 |

Numbers in parentheses represent the value for the highest resolution shell.

^a^ Rmerge = ∑|Ii - Im|/∑Ii, where Ii is the intensity of the measured reflection and Im is the mean intensity of all symmetry related reflections.

^b^ Rwork = Σ||Fobs| - |Fcalc||/Σ|Fobs|, where Fobs and Fcalc are observed and calculated structure factors.

^c^ Rfree = ΣT||Fobs| - |Fcalc||/ΣT|Fobs|, where T is a test data set of about 5% of the total reflections randomly chosen and set aside prior to refinement.
